# Taxane-Induced Conformational Changes in the Microtubule Lattice Activate GEF-H1-Dependent RhoA Signaling

**DOI:** 10.64898/2025.12.20.695343

**Authors:** Joyce C. M. Meiring, Varsha Mahapatra, Molly S.C. Gravett, Daan Morren, Harriet A.J. Saunders, Sung Ryul Choi, Andressa Pelster José, Adela Karhanova, Ioanna Metallidou, Alex Moore, Marèl F.M. Spoelstra, Ruijie Liu, Matteo Giono, Saishree S. Iyer, J. Fernando Díaz, Zdenek Lansky, Kelly E. Stecker, Michel O. Steinmetz, Stuart C. Howes, Lukas C. Kapitein, Anna Akhmanova

**Author notes:** These authors contributed equally (Joyce C. M. Meiring, Varsha Mahapatra).

## Abstract

Taxanes are widely used chemotherapeutic agents that perturb cell division. They also exert effects during interphase, but the underlying mechanisms are poorly understood. Here, we show that taxanes activate RhoA signaling and induce actin remodeling by displacing the RhoA activator GEF-H1 from microtubules. This taxane-induced release of GEF-H1 occurs rapidly, is independent of tubulin post-translational modifications, and can be recapitulated using purified proteins. *In vitro* reconstitution assays combined with analyses of microtubule structure revealed that microtubule binding by GEF-H1 is inhibited by microtubule-stabilizing agents that expand the microtubule lattice, such as taxanes and GMPCPP, but not by others, including GTPγS and discodermolide, which stabilize a compacted microtubule lattice. Our findings demonstrate that alterations in microtubule lattice conformation can activate key signaling pathways, offering new insights into the mode of action of taxanes and the possible origins of their side effects.

## Introduction

Microtubule-targeting agents (MTAs) are widely used to treat different cancers (Dumontet and Jordan, 2010). MTAs are divided into microtubule-destabilizing agents, including colchicine and vinca alkaloids, and microtubule-stabilizing agents, such as taxanes (Florian and Mitchison, 2016; Perez, 2009; Steinmetz and Prota, 2018). Both types of compounds strongly perturb cell division, and this has long been regarded as the explanation of their antitumor effects. However, drugs specifically targeting the formation of the mitotic spindle, such as kinesin-5 inhibitors, showed disappointing results in the clinic, suggesting that interphase effects of MTAs likely contribute to their clinical efficacy (Komlodi-Pasztor et al., 2012; Shi and Mitchison, 2017). Agents that target mitosis specifically are thought to have little success due to their narrow therapeutic window, combined with a slow doubling time of tumors, typically no faster than 1-2 months (Tubiana, 1989; Yan et al., 2020). Moreover, there are direct indications that interphase pathways induced by stabilizing or destabilizing MTAs may promote cancer cell death (Janssen et al., 2013; Mertens et al., 2023; Vennin et al., 2023), but the underlying molecular mechanisms are currently unknown.

The major interphase effect of microtubule-destabilizing agents is the activation of signaling by the small GTPase RhoA (Horin et al., 2025). This effect depends on the RhoA activator guanine nucleotide exchange factor H1 (GEF-H1), which is sequestered on microtubules in inactive form, but becomes active upon release from microtubules (Azoitei et al., 2019; Birkenfeld et al., 2008; Kashyap et al., 2019; Krendel et al., 2002). Since taxanes stabilize microtubules, they would not be expected to affect GEF-H1 activity. However, microtubule stabilization can potentially affect microtubule-associated components of signaling cascades by altering their turnover and tubulin post-translational modifications (Magiera et al., 2018). Previous studies suggested that GEF-H1 avoids acetylated microtubules (Deb Roy et al., 2024; Seetharaman et al., 2022). The underlying mechanism is likely to be indirect, because GEF-H1 binds to the outer microtubule surface (Choi et al., 2025), whereas acetylation occurs on the luminal microtubule surface (Howes et al., 2014; Iuzzolino et al., 2024; Luo et al., 2025). Acetylation typically occurs on stable microtubules (Cambray-Deakin and Burgoyne, 1987; Piperno et al., 1987), and stable microtubules were recently found to have a distinct, expanded conformation (de Jager et al., 2025). This conformation, characterized by an increased longitudinal spacing between tubulin dimers, is also induced by taxanes (Alushin et al., 2014; de Jager et al., 2025; Kellogg et al., 2017). Hence, we set out to investigate whether taxanes can trigger GEF-H1 mediated RhoA activation through direct modification of the microtubule lattice. Using a combination of cell biology and *in vitro* reconstitution assays, we show that taxanes indeed can rapidly activate RhoA by displacing GEF-H1 from microtubules through a direct mechanism that involves an alteration of microtubule lattice conformation.

## Results

### Taxanes activate RhoA through GEF-H1

To measure RhoA activation, we used HT1080 fibrosarcoma cells stably expressing the RhoA2G FRET sensor (Fritz et al., 2013). For better reproducibility, we serum starved cells to lower baseline RhoA activation, since lysophosphatidic acid in serum is a potent inducer of RhoA signaling (Ridley and Hall, 1992). As expected, RhoA activity was low in serum-starved cells, but showed a rapid transient increase with serum addition (Fig.1A, B). The microtubule destabilizing agent nocodazole, which was used as a positive control for GEF-H1 activation, caused complete microtubule disassembly and induced a slower, but persistent increase of RhoA activity (Fig.1A, B). A similar, although weaker RhoA activation was also triggered by the addition of a high concentration (10 µM) of paclitaxel (Fig. 1A, B). The effects of both nocodazole and paclitaxel, but not the effect of serum on RhoA activity were largely suppressed in GEF-H1 knockout cells (Fig. 1C, D, Fig. S1A-D), indicating that both MTAs activate RhoA through GEF-H1.

**Figure 1.**
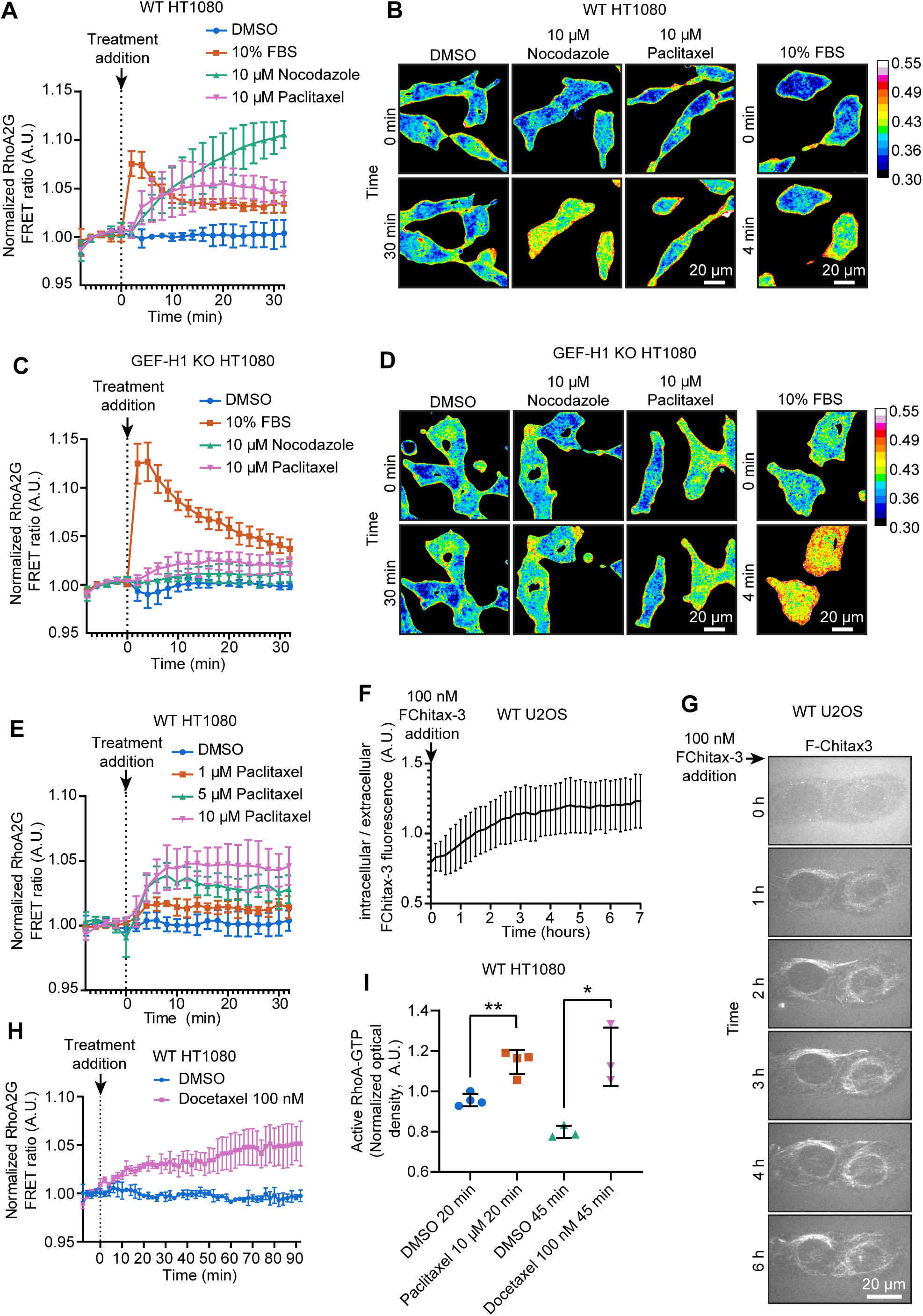
Paclitaxel activates RhoA in cells in a GEF-H1 dependent manner. (A-D) Serum starved HT1080 wild type (WT) (A, B) or GEF-H1 knockout (KO) (C, D) cells stably expressing RhoA2G FRET sensor were first imaged live for 8 minutes without treatment to establish a baseline, then treated with either paclitaxel, nocodazole, serum (10% Fetal Calf Serum) as a positive control or dimethyl sulfoxide (DMSO) as a solvent control at time = 0 min and imaged for another 32 minutes. (A, C) Graphs show mean ± standard deviation of the experimental means of 3 independent experiments. Data were normalized to the average baseline collected before treatment. (B, D) Representative stills of RhoA2G FRET ratio of cells before (time = 0 min) and after treatment (time = 30 min or 4 min). RhoA2G FRET sensor fluorescence was used to create masks of the cells, and all signals outside of the mask were cleared. (E, H) Serum starved HT1080 wild type (WT) cells stably expressing RhoA2G FRET sensor were first imaged live for 8 minutes without treatment to establish a baseline, then treated with either 1, 5 or 10 µM paclitaxel or 100 nM docetaxel or DMSO as a solvent control and imaged for a further (E) 32 minutes or (H) 92 minutes. FRET ratios were normalized to the average of the baseline collected before treatment addition. Plots show the mean ± standard deviation of the experimental means from 3 experiments. (F, G) U2OS cells were imaged on a spinning disc microscope from the moment fluorescent taxane FChitax-3 was added to the cell medium for 7 hours. (F) Plot shows mean ± standard deviation of ratio of intracellular / extracellular FChitax-3 fluorescence (n = 5, from one representative experiment). (G) Representative frames showing FChitax-3 accumulating on microtubules in cells. (I) Serum starved HT1080 were treated with DMSO or MTAs as shown for 20 or 45 min before harvesting lysates and quantifying active RhoA-GTP via a RhoA G-LISA assay; plots show the mean ± standard deviation of the experimental means (DMSO and paclitaxel 20 min, n = 4 experiments, DMSO and docetaxel 45 min, n = 3 experiments). A Welch’s t-test was used to test for statistical significance, * = p<0.05, ** = p<0.01.

In short-term (30 min) assays, paclitaxel-induced activation of RhoA was weaker than that triggered by nocodazole. At least 5 µM concentration of the compound was needed to induce a substantial effect (Fig. 1A, E), whereas taxane concentrations in plasma of cancer patients during chemotherapy reach a concentration of 0.1-1 µM (Brouwer et al., 2000; Gardner et al., 2008). Importantly, docetaxel, a taxane with a higher affinity to microtubules (Buey et al., 2005), which is frequently used for cancer chemotherapy (Imran et al., 2020), could activate RhoA signaling already at a concentration of 100 nM (Fig. 1H). The magnitude of RhoA activation by docetaxel increased over time (Fig. 1H), likely due to a slow but persistent accumulation of the drug, an idea supported by 7-hour long live imaging of cell accumulation of a fluorescent taxane, Fchitax-3 (Fig. 1F, G). RhoA activation with 10 µM paclitaxel or 100 nM docetaxel was also confirmed using a RhoA G-LISA assay (Fig. 1I). These results demonstrate that taxanes can trigger GEF-H1 mediated RhoA activation at clinically relevant concentrations.

### Paclitaxel causes GEF-H1-dependent actin remodeling

The main downstream effect of RhoA activation is the stabilization of filamentous actin and increased actomyosin contractility, typically visualized as an increase in stress fiber formation in 2D cell cultures. Serum-starved HT1080 cells were elongated and had only a few stress fibers and lamellipodia at the leading edge (Fig. 2A). Treatment with Y-27632, an inhibitor of Rho-associated protein kinase (ROCK), a kinase acting downstream of RhoA (Amano et al., 2010; Lawson and Ridley, 2018; Riento and Ridley, 2003), had no strong effect on actin organization or cell shape in these conditions (Fig. 2A, C, E), in agreement with serum starvation reducing baseline RhoA activity. In contrast, microtubule disassembly with nocodazole induced formation of stress fibers, blebs and cell rounding (Fig. 2A, C, E). A very similar effect was observed when microtubules were stabilized with paclitaxel (Fig. 2A, C, E). However, in GEF-H1 knockout cells, MTA treatments had no pronounced effect on the actin cytoskeleton, and cells retained their elongated shape (Fig. 2B, D, F). In line with the changes in the actin organization, paclitaxel also increased the abundance of phosphorylated myosin light chain (p-MLC), a downstream target of RhoA signaling (Lawson and Ridley, 2018) (Fig. 2G, H). Taken together, these data show that, similar to microtubule-depolymerizing agents, paclitaxel triggers GEF-H1-dependent actin reorganization.

**Figure 2.**
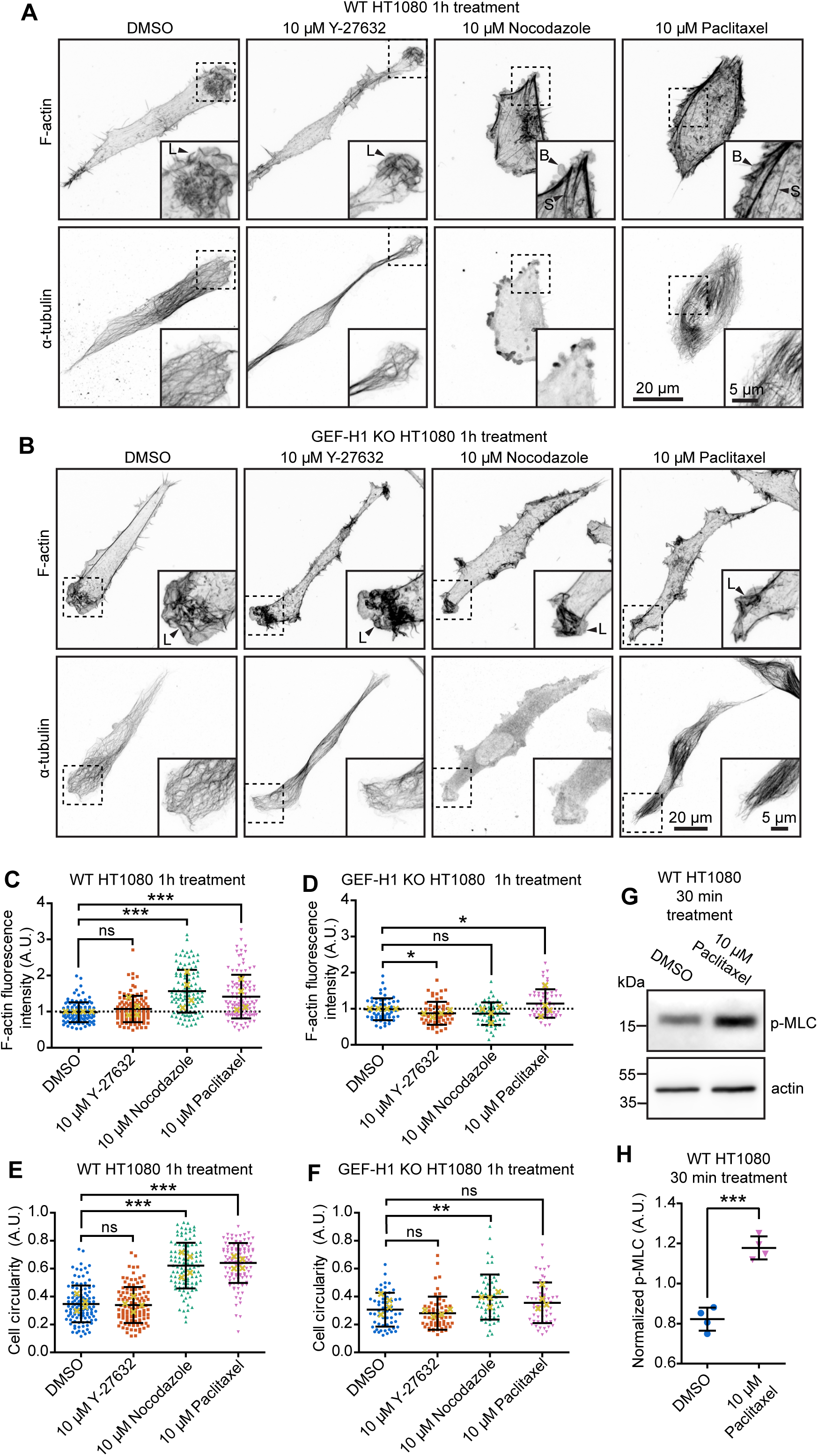
Paclitaxel activates downstream effectors of RhoA in a GEF-H1-dependent manner. (A-F) Serum starved HT1080 WT (A, C, E) or GEF-H1 KO (B, D, F) cells were treated with paclitaxel, nocodazole, Y-27632 (ROCK inhibitor) or DMSO as a solvent control for 1 hour before fixing and staining for F-actin and α-tubulin. (A, B) Representative images, where dashed box shows region highlighted in zoom panel; examples of lamellipodia (L), blebs (B) and stress fibers (S) are highlighted with labelled arrows. (C-F) Plots show F-actin staining intensity (C, D) or cell circularity (E, F) normalized to experimental mean of DMSO control; individual cells are shown as colored symbols, yellow crosses show experimental means, brackets show standard deviation based on individual cell values. For (C, E) n = 112 cells from 4 experiments; for (D, F) n = 58 cells from 4 experiments. (G, H) Serum starved HT1080 WT cells treated with paclitaxel or DMSO as a solvent control for 30 minutes before harvesting cell lysates and blotting for p-MLC and actin as a loading control. (G) Representative blot from 1 experimental replicate. (H) Plot shows analyzed p-MLC from Western blot normalized to actin loading control and experimental mean, brackets show standard deviation, n = 4 experiments. Mann-Whitney test was used to analyze statistical significance for F-actin fluorescence and cell circularity quantifications, Welch’s t-test was used to analyze p-MLC quantifications, ns = non-significant, * = p<0.05, ** = p<0.01, *** = p<0.001.

### Paclitaxel displaces GEF-H1 from microtubules in cells

Since microtubule-destabilizing agents activate GEF-H1 by displacing it from microtubules (Azoitei et al., 2019; Birkenfeld et al., 2008; Kashyap et al., 2019; Krendel et al., 2002), we next investigated whether paclitaxel could also release GEF-H1 from microtubules. GEF-H1 immunostaining in cells shows strong variability and ambiguous microtubule co-localization, making reliable quantitative analysis difficult (Fig. S1D). To circumvent this problem, we overexpressed GFP-tagged full length GEF-H1 (GFP-GEF-H1) at low levels, and quantified GFP signal to ensure comparable expression levels between treatments (Fig. 3A-E). We chose U2OS for these experiments as these are more strongly adherent than HT1080, and therefore less prone to rounding up in response to treatment with taxanes. As expected, GFP-GEF-H1 bound along most microtubules in U2OS cells but was barely detected on acetylated microtubules (Fig.3A yellow dotted line, Fig. 3C), in line with previous studies (Deb Roy et al., 2024; Seetharaman et al., 2022). Treatment of cells with 10 µM paclitaxel resulted in dissociation of GEF-H1 from microtubules (Fig. 3A). This effect could be observed and quantified in fixed cells (Fig. 3A, D) and directly visualized by live imaging (Fig. 3F, Movie 1). GFP-GEF-H1 displacement was not simply a result of suppression of microtubule growth, because another microtubule-stabilizing agent binding at the taxane-site, discodermolide (Buey et al., 2005; Longley et al., 1991), which also efficiently inhibits microtubule elongation detected by staining EB1/EB3 positive microtubule plus ends (Fig. S2A), had no effect on GFP-GEF-H1 binding to cellular microtubules (Fig. 3A, D, G, Movie 2). To narrow down the origin of GEF-H1’s sensitivity for taxanes, we looked at the microtubule binding domain of GEF-H1, which consists of a zinc-finger-like C1 domain flanked by intrinsically disordered regions (amino acids 1-136; Fig. S1A). Since full length GEF-H1 is a coiled-coil dimer (Choi et al., 2025), we artificially dimerized the shorter fragment using a GCN4 (G4) leucine zipper. The GFP-GEF-H1_1-136_-G4 was also displaced from microtubules by paclitaxel but not by discodermolide (Fig. S2B). This suggests that the first 135 amino acids of GEF-H1 help confer its sensitivity to taxane-induced lattice conformations. We also tested shorter fragments of GEF-H1, such as the dimeric version of C1 domain alone (amino acids 28-100) dimerized using GCN4, but with this construct, we only detected very weak enrichment on microtubules, while versions without the GCN4 did not show any detectable microtubule binding.

**Figure 3.**
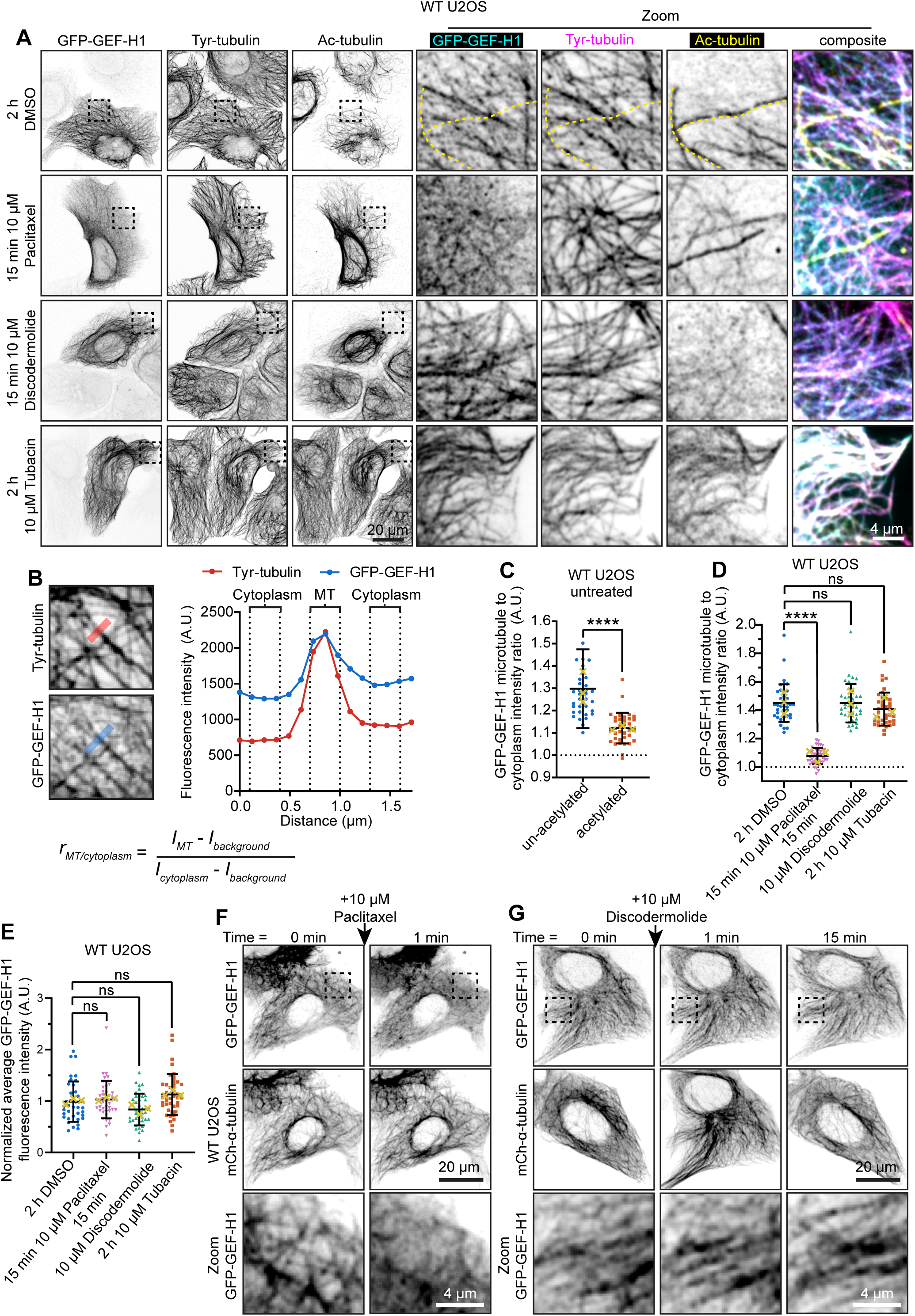
Paclitaxel displaces GEF-H1 from microtubules in cells. (A-E) U2OS cells transfected with GFP-GEF-H1 at low levels were treated for 15 minutes with paclitaxel or discodermolide or treated for 2 hours with Tubacin (inhibitor of tubulin deacetylase HDAC6) or DMSO solvent control before fixing and staining for tyrosinated α-tubulin (tyr-tubulin) or acetylated α-tubulin (ac-tubulin). (A) Representative images, where dashed boxes indicate regions enlarged in zoom panels, and dashed yellow line highlights acetylated microtubules devoid of GFP-GEF-H1. (B) GFP-GEF-H1 on microtubules was analyzed by first drawing line scans perpendicularly across microtubules flanked by cytoplasm and using the maxima of the tyr-tubulin staining to define microtubule and cytoplasmic regions within GFP-GEF-H1 intensity values. The ratio of GFP-GEF-H1 on a microtubule was subsequently calculated by dividing the intensity on the microtubule by the intensity in the cytoplasm. (C) Plot of GFP-GEF-H1 microtubule to cytoplasm ratios of acetylated vs un-acetylated microtubules in the same cell (n = 35, from 3 experiments). (D-E) (D) Plot of GFP-GEF-H1 microtubule to cytoplasm intensity ratios, (E) Plot of GFP-GEF-H1 intensity normalized to experimental mean (DMSO n = 37; paclitaxel n = 41; discodermolide n = 37; Tubacin n = 39; from 3 experiments). (C-E) Individual cell means are shown as colored symbols, yellow crosses show experimental means, and brackets show standard deviation based on individual cell values. A Wilcoxon matched-pairs signed rank test was used to test for statistical significance of GFP-GEF-H1 microtubule binding between acetylated and non-acetylated microtubules and a Welch’s t-test was used to compare between DMSO and drug treated cells, ns = not significant, **** = p<0.0001. (F, G) U2OS cells were transfected with mCherry-α-tubulin and GFP-GEF-H1 and imaged live on a spinning disc microscope before (Time = 0 min) and after (Time = 1 min, 15 min) addition of paclitaxel (F) or discodermolide (G) as a microtubule stabilizing control that does not expand microtubule lattice.

GEF-H1 dissociation from microtubules in paclitaxel-treated cells occurred within 1 minute of drug addition (Fig. 3F, Movie 1), whereas discodermolide treatment had no effect (Fig. 3G, Movie 2). One minute is insufficient to trigger marked microtubule acetylation, since even cells treated with paclitaxel for 15 minutes did not show significant over-acetylation of microtubules (Fig. 3A). Moreover, treatment of cells with a tubulin deacetylase HDAC6 inhibitor Tubacin (Haggarty et al., 2003) or overexpression of the tubulin acetylase αTAT1 (Akella et al., 2010), both of which caused acetylation of all cellular microtubules, did not inhibit the binding of GEF-H1 to microtubules (Fig. 3A, D; Fig. S2C,F). In fact, we saw a subtle increase in GEF-H1 binding to microtubules upon αTAT1 overexpression (Fig. S2F), despite there being no difference in GFP-GEF-H1 fluorescence intensities (Fig. S2G). Induction of microtubule de-tyrosination (removal of the C-terminal tyrosine of α-tubulin), another modification associated with microtubule stability (Magiera et al., 2018), by overexpressing detyrosinating enzymes VASH2 + SVBP and MATCAP (Aillaud et al., 2017; Landskron et al., 2022; Nieuwenhuis et al., 2017), also either did not impact the localization of GEF-H1 to microtubules or showed a slight increase in binding (Fig. S2D-F). Taken together, our data indicate that paclitaxel displaces GEF-H1 from microtubules through a rapid mechanism that cannot be explained by altered microtubule dynamics or tubulin post-translational modifications.

### Characterization of GEF-H1-microtubule interactions by *in vitro* reconstitution assays

Based on cellular data, we hypothesized that GEF-H1 binding to microtubules is sensitive to changes in lattice conformation and set out to test this idea using assays with purified proteins. Dynamic microtubules were grown in the presence of fluorescently labeled tubulin from microtubule seeds stabilized with the slowly hydrolysable GTP analog GMPCPP and observed by Total Internal Reflection Fluorescence (TIRF) microscopy (Fig. 4A) (Bieling et al., 2007; Saunders et al., 2025). GFP-GEF-H1 and GFP-GEF-H1_1-136_-G4 were purified from transiently transfected HEK293T cells (Fig. S3A, B). To evaluate protein purity, we used mass spectrometry and detected only minor contamination of the GFP-GEF-H1 preparation with HSP70 chaperones and 14-3-3 proteins (Fig. S3C). Both GFP-GEF-H1 and GFP-GEF-H1_1-136_-G4 efficiently accumulated on dynamic microtubule lattices already at very low concentrations (500 pM for the full-length GEF-H1 and 3 nM for GEF-H1_1-136_-G4) but strongly avoided GMPCPP-stabilized seeds (Fig. 4B-E). We used Interference Reflection Microscopy (IRM) to confirm that GMPCPP-stabilized seeds and the dynamic lattices grown from these seeds consist of 14 protofilaments, as published previously (Hyman et al., 1995; Rai et al., 2021) (Fig. S3D, E). Since in mammalian cells the majority of microtubules consist of 13 protofilaments (Chaaban and Brouhard, 2017), these data suggest that GEF-H1-microtubule interaction is not particularly sensitive to protofilament number. Furthermore, avoidance of GMPCPP seeds could not be due to the lack of GTP hydrolysis by β-tubulin within the seed, because 500 pM GFP-GEF-H1 efficiently decorated microtubules polymerized in the presence of non-hydrolysable GTP analog GTPγS (Fig. 4F).

**Figure 4.**
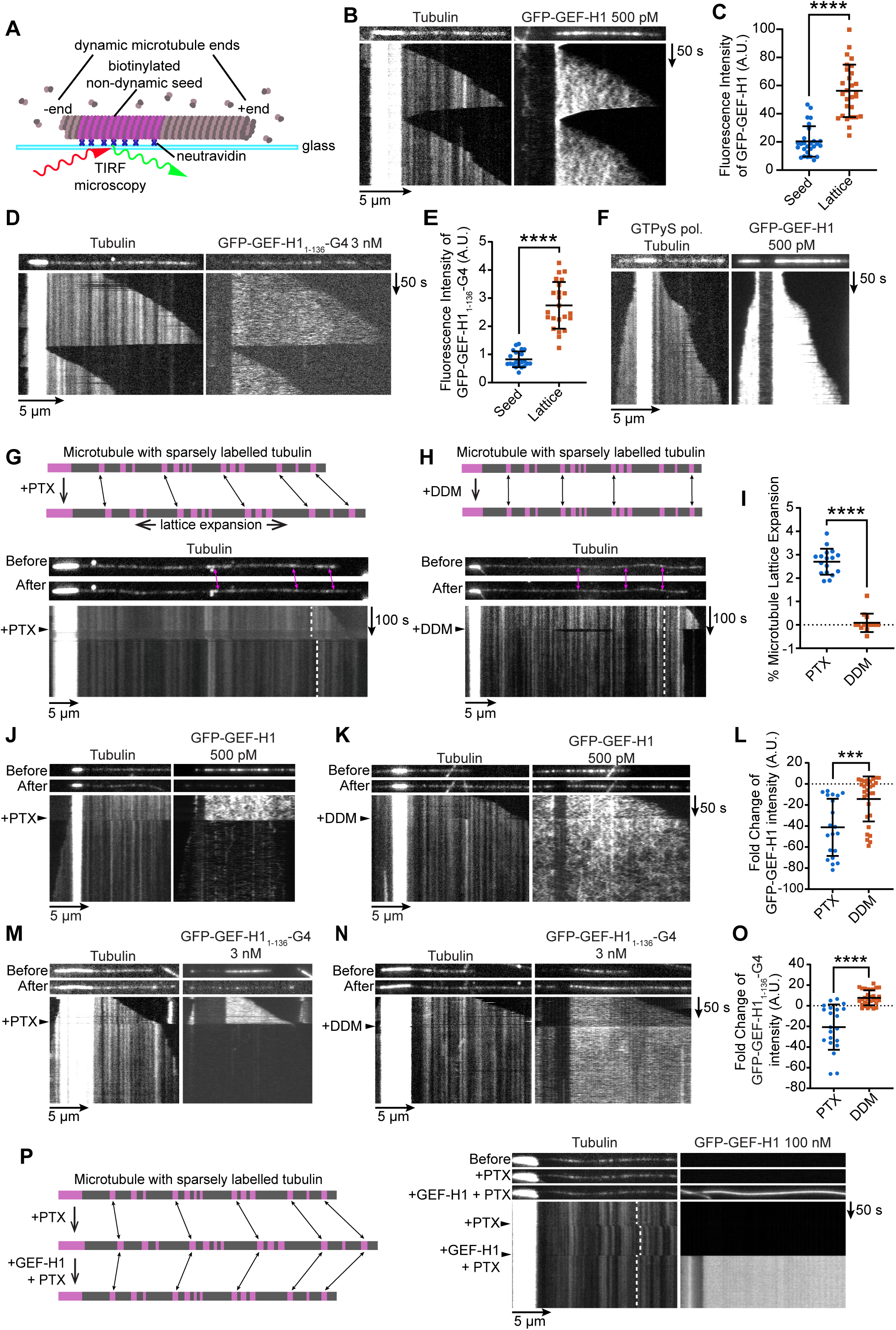
Characterization of GEF-H1-microtubule interactions by *in vitro* reconstitution assays. (A) Scheme of the *in vitro* assay with dynamic microtubules. (B,D) Still images and kymographs of representative microtubules grown with 500 pM GFP-GEF-H1 (B) or 3nM GFP-GEF-H1_1-136_-G4 (D). Tubulin channel shows bright GMPCPP seeds (left); GFP-GEF-H1 channel (right) shows binding to the dynamic GDP lattice but exclusion from GMPCPP seeds. (C,E) Plots showing mean ± standard deviation of intensity of (C) GFP-GEF-H1 (seed n = 27, lattice n = 28; from 3 experiments) and (E) GFP-GEF-H1_1-136_-G4 (seed n = 22, lattice n = 23, from 3 experiments) quantified at GMPCPP seed vs GDP lattice. (F) Still images and kymographs of a representative microtubule grown with 500 pM GEF-H1 using GTPγS instead of GTP show that GEF-H1 still avoids GMPCPP seeds but binds the GTPγS lattice. (G, H, top) Schemes of sparsely labelled microtubules used to detect microtubule expansion (or lack thereof) upon wash-ins. (G, H, bottom) Still images of representative microtubules before and after wash-ins, illustrating speckle shift upon the addition of paclitaxel (PTX, G) but not discodermolide (DDM, H). The frame of the wash-in is indicated in the kymographs with an arrowhead; dotted lines highlight speckle shift or lack thereof. (I) Plot showing percentage of expansion with PTX or DDM (PTX n = 18, DDM n = 18; from 3 experiments). (J-O; Top): Still images of representative microtubules before and after PTX or DDM wash-in in assays with microtubules grown in the presence of (J, M) 500pM GFP-GEF-H1 or (K, N) 3nM GFP-GEF-H1_1-136_-G4; tubulin (left) and GFP channels (right) are shown. Bottom panels: corresponding kymographs, with wash-in frame indicated with an arrowhead. (L, O) Plots showing fold change in (L) GFP-GEF-H1 (PTX n = 20, DDM n = 29; from 3 experiments) or (O) GFP-GEF-H1_1–135_-G4 (PTX n = 21, DDM n = 31; from 3 experiments) intensity upon PTX or DDM wash-in. (P, left) Scheme of the experiment used to detect compaction by GEF-H1. (P, right). Still images before, after PTX, and after 100 nM GEF-H1 addition; arrowheads indicate the frames of wash-ins in the kymographs; dotted lines indicate speckle shift. Mann-Whitney test was used to test for statistical significance, *** = p<0.001,**** = p<0.0001.

To test if microtubule-stabilizing agents directly affect GEF-H1 binding, we performed wash-in assays with 20 µM paclitaxel or 200 nM discodermolide. Discodermolide has a significantly higher affinity to microtubules than paclitaxel and is a known hypernucleator of microtubules *in vitro* (Hung et al., 1996; ter Haar et al., 1996). Hence, to avoid spontaneous nucleation interfering with our assays, we used a lower concentration which was still able to prevent depolymerization *in vitro*. Additionally, to prevent spontaneous microtubule nucleation, the concentration of soluble tubulin during wash-in was reduced from 17 µM to 5 µM. Paclitaxel is known to expand the microtubule lattice in the longitudinal direction (Alushin et al., 2014; Kellogg et al., 2017), an effect that could be readily observed by imaging microtubules sparsely labelled by fluorescent tubulin (Fig. 4G, I). In contrast, discodermolide did not expand the microtubule lattice (Fig. 4H, I). To strengthen this conclusion, we determined the lattice parameters of 13 pf and 14 pf microtubules stabilized with discodermolide using cryo-electron microscopy (cryo-EM) followed by single particle analysis. We found that discodermolide-stabilized microtubules indeed had a compacted lattice, and had lattice parameters that were highly similar to those of compacted GTPγS-bound lattices (Fig. S4A,B; 81.5Å dimer rise for 13 pf and 14 pf discodermolide-stabilized microtubules, compared to 81.9 Å and 81.7Å for 13 pf and 14 pf microtubules assembled with GTPγS (Zhang et al., 2018)). The binding of GFP-GEF-H1 and GFP-GEF-H1_1-136_-G4 to microtubules was not altered by discodermolide addition, but both proteins were very rapidly displaced after paclitaxel wash-in (Fig. 4J-O). Furthermore, the addition of a 200 fold higher concentration (100 nM) of GFP-GEF-H1 rapidly compacted microtubules that were expanded by paclitaxel addition (Fig. 4P). These data strongly support the notion that GEF-H1 binding to microtubules is very sensitive to microtubule lattice conformation: it is favored by compacted, GDP-, GTPγS- or discodermolide bound lattices but is incompatible with GMPCPP- or paclitaxel-stabilized, expanded lattice conformations. However, at sufficiently high concentrations, GEF-H1 can impose its preferred compacted lattice state even on microtubules expanded by paclitaxel.

## Discussion

In this study, we have demonstrated that taxanes can rapidly stimulate RhoA signaling by displacing the RhoA activator GEF-H1 from microtubules, thereby triggering actomyosin contractility and cell rounding. By combining experiments in cells with *in vitro* reconstitutions using purified components, we revealed that GEF-H1 detachment is driven by taxane-induced conformational changes in the microtubule lattice. Next to microtubule post-translational modifications and tubulin isotypes, lattice conformation is emerging as an important, understudied constituent of the “tubulin code”, the biochemical and functional diversification of microtubule subsets in the same cell (Akhmanova and Kapitein, 2022; Janke and Magiera, 2020; Verhey and Ohi, 2023).

Diversity of microtubule conformations includes differences in protofilament number, longitudinal compaction or expansion, and skew (Chaaban and Brouhard, 2017; Kellogg et al., 2017; LaFrance et al., 2022; Paquette et al., 2025) (Fig. S4C). Our data indicate that GEF-H1 can bind to both 13- and 14-protofilament microtubules, and the effect of paclitaxel on GEF-H1 binding is too fast to be caused by a switch in protofilament number. 13- and 14-protofilament microtubules have very different skews (Fig. S4B), indicating that GEF-H1 may not be sensitive to this lattice parameter. Since GEF-H1 avoids expanded GMPCPP- or paclitaxel-stabilized microtubule lattices but binds well to compacted GDP-, discodermolide- or GTPyS-bound lattices, our data point to GEF-H1 sensitivity to longitudinal expansion/compaction, which mostly affects the interface around β-tubulin E-site (Fig. S4C) (Alushin et al., 2014). However, recent cryo-EM structural studies have shown that the C1 domain of GEF-H1 (amino acids 28–100) binds 13-protofilament paclitaxel-stabilized microtubules at the junction between two neighboring tubulin dimers (Choi et al., 2025), which is distant from the β-tubulin E-site (Fig. S1A and S4D). Interestingly, microtubules in the complex with C1 domain displayed a partially compacted lattice and a protofilament skew similar to that observed in GTPγS- and discodermolide-stabilized microtubules (Fig. S4A,B). These data suggest that, under saturating concentrations, the C1 domain alone can alter microtubule lattice conformation through allosteric effects that are initiated at the GEF-H1 C1 binding site and are transmitted to the β-tubulin E-site. The exact nature of these allosteric effects is currently unclear. More work using longer GEF-H1 fragments with higher microtubule-binding affinity and microtubule lattices to which GEF-H1 binds well, such as GTPγS-stabilized microtubules, might help to understand the structural basis of GEF-H1 microtubule conformation preferences.

Our cellular, *in vitro* and cryo-EM work established that discodermolide, a compound that stabilizes microtubules and perturbs cell division similar to taxanes (ter Haar et al., 1996), is a very useful control for taxanes. Cryo-EM data showed that unlike taxanes, discodermolide promotes the formation of compacted microtubule lattices; this is in line with recent analysis based on fiber diffraction (Lucena-Agell et al., 2025). Similar to paclitaxel, discodermolide inhibited microtubule growth in cells but did not trigger GEF-H1 detachment from microtubules. This control will be valuable for distinguishing the impact of changes in lattice conformation from effects on microtubule growth and mitotic arrest in future investigations into the mechanism of action of microtubule-stabilizing agents. Additionally, discodermolide can be utilized to distinguish the sensitivity of microtubule binding proteins to lattice conformation versus microtubule dynamics. Our study hence provides a useful framework for elucidating the sensitivity of microtubule binding proteins and cellular processes to the conformational tubulin code.

An important feature of microtubule lattice plasticity, in contrast to biochemical modifications, is that proteins serving as “conformational code readers” can become “code writers” at a high concentration (Siahaan et al., 2022). This explains why the sensitivity of GEF-H1 to taxanes has been overlooked previously (Kashyap et al., 2019): when strongly overexpressed, GEF-H1 can induce its preferred lattice conformation that cannot be altered by paclitaxel. However, endogenous GEF-H1 is a signaling molecule present at levels that are sufficient to detect but not to alter microtubule lattice compaction. GEF-H1 avoids acetylated, stable microtubules, likely because their lattices are expanded (de Jager et al., 2025; Jansen et al., 2023) and thus more similar in conformation to paclitaxel- and GMPCPP-stabilized microtubules. This might prevent, for example, sequestration of GEF-H1 on stable perinuclear microtubules, so that it can control RhoA activity in response to local changes in the density of dynamic microtubules near the cell margin (Azoitei et al., 2019).

Activation of RhoA by taxanes was likely overlooked previously because in cycling cultured cells, this effect is quickly overshadowed by mitotic arrest, especially at low concentrations, when taxanes accumulation in cells is slow. The ability of taxanes to activate RhoA can be relevant for understanding their efficacy in cancer therapy, because RhoA can promote anti-tumor immune responses (Bros et al., 2019; Kalim et al., 2021; Kashyap et al., 2019). RhoA signaling also might potentially contribute to cancer cell death, because it can promote apoptosis in low adhesion environments by increasing cell contractility, explaining why ROCK inhibitors are a standard component of media for stem cell and organoid isolation and maintenance in culture (Koyanagi et al., 2008; Martin-Ibanez et al., 2008; Minambres et al., 2006; Sato et al., 2009; Walker et al., 2010). On the other hand, misregulation of RhoA might contribute to the side effects of taxanes, such as neuropathy (McCray et al., 2021; Ohsawa et al., 2011). On a broader scale, taxane-induced changes of microtubule lattice conformation may alter binding of additional microtubule-associated proteins and thus perturb other signaling pathways besides RhoA - a concept deserving future study.

## Acknowledgements

We thank S. Etienne-Manneville (Institut Pasteur, Paris, France), M. Martin (University of Brussels, Belgium), T. Brummelkamp (Netherlands Cancer Institute, Amsterdam, the Netherlands), D. Trono (EPFL, Lausanne, Switzerland), and F. Zhang (Broad Institute of MIT and Harvard, Cambridge, USA) for the gift of materials. This work was supported by the Netherlands Organization for Scientific Research (NWO) Gravitation programme IMAGINE! (project number 24.005.009), ENW-XL grant (project number OCENW.XL21.XL21.048), the European Research Council Synergy grants PushingCell (project number 101071793) and TUBULINCODE (project number 101071583), the European Union’s Horizon Europe research and innovation programme under the Marie Skłodowska-Curie grant agreement No. 101153345, institutional support from the CAS (RVO: 86652036), a grant from the Swiss National Science Foundation (31003A_166608; to M.O.S.), the Imaging Methods Core Facility at BIOCEV supported by the MEYS CR (Large RI Project LM2018129 Czech-BioImaging) and ERDF (project no. CZ.02.1.01/0.0/ 0.0/16_013/0001775), Ministerio de Ciencia e Innovación (MICIN) and FEDER PID2022−136765OB-I00, GAUK (project number 204125), and the NWO-funded Netherlands Proteomics Center through the National Roadmap for Large-scale Infrastructures program XOmics (Project 184.034.019).

## Author Contributions

J.C.M.M. and V.M. performed experiments, analyzed data and wrote the paper, M.S.C.G. and A.P.J. performed and analyzed cryo-electron microscopy experiments with discodermolide-stabilized microtubules, D.M., H.A.J.S., A.K. and M.F.M.S performed experiments and analyzed data, I.M. performed experiments, M.G. generated recombinant constructs, A.M., R.R.L and S.R.C. analyzed data, S.S.I. purified protein, J.F.D. contributed reagents and acquired funding, K.E.S. supervised and analyzed mass spectrometry experiments, Z.L. supervised IRM experiments, M.O.S supervised structural analyses and acquired funding, S.H. supervised and analyzed cryo-electron microscopy experiments and structural analyses, acquired funding and wrote the paper, L.C.K. supervised *in vitro* experiments, acquired funding and wrote the paper, A.A. coordinated the project, acquired funding and wrote the paper. All authors reviewed and edited the paper.

## Competing financial interests

The authors declare no competing financial interests.

## Supplemental Videos

**Video 1. Paclitaxel addition displaces GFP-GEF-H1 from microtubules in cells.**

U2OS cells transfected with GFP-GEF-H1 (left) and mCherry-α-tubulin (mCh-α-tubulin; right) were imaged for 10 minutes before the addition of 10 µM paclitaxel, indicated by orange border and “+PTX”, and subsequently imaged for another 20 minutes.

Video 2. Discodermolide addition does not displace GFP-GEF-H1 from microtubules in cells.

U2OS cells transfected with GFP-GEF-H1 (left) and mCherry-α-tubulin (mCh-α-tubulin; right) were imaged for 10 minutes before the addition of 10 µM discodermolide, indicated by blue border and “+DDM”, and subsequently imaged for another 20 minutes.

## Methods Cell culture

HT1080, U2OS and HEK293T cells were cultured in Dulbecco’s Modified Eagle Medium (DMEM; Capricorn Scientific) supplemented with 10% Fetal Bovine Serum (FBS; Corning) and 100U/mL Penicillin and 100 µg/mL Streptomycin (1% Pen Strep; Sigma). Cells were kept at 37 °C with 5% CO2 and routinely tested for mycoplasma using a commercial assay (Mycoaltert assay, LT07-518; Lonza).

## Serum starvation

For assays directly or indirectly assessing RhoA activation, cells were first serum-starved to lower baseline RhoA activity as follows: cells were plated at 10-15% confluency in normal culture medium containing 10% FBS, transferred to medium containing 0.5% FBS 24 h after plating and changed to FBS free medium another 24 h after that and incubated overnight (16-18 h) before commencing the experiment.

## DNA constructs

GFP-GEF-H1 was a gift from S. Etienne-Manneville (Institut Pasteur, Paris, France). For protein purification, a strepII tagged GFP-GEF-H1 was generated by using Gibson assembly using GFP-GEF-H1 as a template to clone into a SII-GFP expression vector. For GFP- GEF-H1_28-100_, GFP-GEF-H1_28-100_-G4, GFP-GEF-H1_1-136_ and GFP- GEF-H1_1-136_-G4, a synthetic gene encoding amino acids 1–136 of GEF-H1 (UniProt Q92974-1), comprising the C1 domain and its adjacent disordered regions, was purchased from Genewiz–Azenta. This was used as a template to amplify GEF-H1_28-100_ and GEF-H1_1-136_. Gibson assembly was used to clone both GEF-H1_1-136_ and GEF-H1_28-100_ into either a SII-GFP expression vector or a modified SII-GFP vector containing a GCN4 coiled-coil domain to promote dimerization. mCh-α-tubulin consisted of a human α-tubulin fused with mCherry (van der Vaart et al., 2011). pLenti-RhoA2G (Fritz et al., 2013) was a gift from Olivier Pertz (Addgene #40179), FLAG-aTAT1 (Seetharaman et al., 2022) was a gift from S. Etienne Manneville (Institut Pasteur, Paris, France), VASH2-FLAG and SVBP-FLAG were a gift from M. Martin (University of Brussels, Belgium), and HA-MATCAP (Landskron et al., 2022) was a kind gift from T. Brummelkamp (Netherlands Cancer Institute, Amsterdam, the Netherlands). psPAX2 (Addgene #12260) and pMD2.G (Addgene #12259) lentiviral packaging constructs were a gift from D. Trono (EPFL, Lausanne, Switzerland). pSpCas9(BB)–2A-Puro (PX459) V2.0(Ran et al., 2013) (Addgene #62988) was a gift from F. Zhang (Broad Institute of MIT and Harvard, Cambridge, USA).

## Cell transfection

Cells were plated at 15% confluency the day prior to transfection. A ratio of 1µg plasmid DNA to 3 µL Fugene6 (Promega) transfection reagent was used, transfection mix was prepared in optiMEM (Gibco) according to manufacturer’s instructions. Cells were used for experiments 24 h after transfection.

## Western blotting

Cell lysates were collected on ice using RIPA buffer (20 mM Tris pH 7.5, 100 mM NaCl, 1% NP-40, 0.1% SDS, 0.2% Sodium Deoxycholate, 10 mM EDTA) containing freshly added broad spectrum protease inhibitors (cOmplete protease inhibitor cocktail, Roche, 11873580001) and phosphatase inhibitors (PhosSTOP, Roche, 4906837001), mixed with loading dye (50 mM Tris pH 6.5, 2% SDS, 1.5 mM Bromophenol blue, 8% glycerol) and 5% β-mercaptoethanol, then boiled for 10 minutes. Lysates were run on 5-16% acrylamide gels and transferred, to PVDF membrane (0.45 µm pore size, Merck Millipore, IPVH00010) or nitrocellulose membrane (0.2 µm pore size, Amersham, 10600006) in the case of p-MLC, using wet transfer at 37 V at 4 °C overnight. Membranes were blocked using 5% BSA in Tris buffered saline with 0.1% Tween 20 (TBST) for 2-8 hours then incubated in primary antibody diluted in blocking buffer, rabbit anti-p-MLC (Cell Signaling, 3671S, dil 1:250) or rabbit anti-GEF-H1 (Abcam, ab155785, dil 1:1000) overnight or mouse anti-actin (Sigma, MAB1501, dil 1:4000) for 1 hour. Blots were washed 3 times for 5 min each with TBST after primary antibody incubation, then incubated with goat anti-mouse HRP conjugated secondary (Agilent, P044701-2, 1:2500 dil) or swine anti-rabbit HRP conjugated secondary (Agilent, P039901-2, 1:2500 dil). Blots were again washed 3 times for 5 min each with TBST then developed using ECL Western blotting substrate (Promega, W1015) and imaged using an Amersham ImageQuant 800, taking care not to saturate the signal. Actin was used as a loading control. Quantifications of p-MLC blots were performed in Image J. p-MLC signal was first normalized to the actin loading control, and then to the experiment average.

## GEF-H1 knockout (KO) cell line generation

Anti-GEF-H1 targeting sequence 5’- atgggcacctcttcaccacc -3’ was cloned into the pSpCas9(BB)–2A-Puro (PX459) V2.0 plasmid using the BbsI restriction site. This GEF-H1 targeting CAS9 plasmid was then transfected into HT1080 cells as described above. 24 hours after transfection, 3 μg/mL puromycin (InvivoGen, ant-pr-1) was added to the medium to select for transfected cells for 24 hours. From the surviving cell pool single cells were plated in a 96 well plate and expanded into clonal cell lines. These were screened for GEF-H1 KO by immunofluorescence, Western blotting and by performing targeted sequencing of genomic DNA. Genomic DNA was extracted using the GeneJET Genomic DNA Purification Kit (Thermo Scientific, K0722) then amplified using the following primers: FW 5’-gctctgctttaggaactggtgt-3’ and RV 5’-gagagaagagtgaccctcatgg-3’, then repurified and sent for sequencing using the RV primer.

## RhoA2G stable cell line generation

Lentivirus was used to generate stable cell lines expressing the RhoA2G FRET sensor. Lentivirus was produced in HEK293T cells; for this purpose, cells were plated in a 10 cm dish to be 90% confluent on the day of transfection. Cells were transfected with 15 μg pLenti-RhoA2G, 10 μg psPAX2 and 5 μg pMD2.G using 90 μL MaxPEI (1 mg/mL Polyethylenimine, Polysciences) in optiMEM (1.2 mL total volume). Transfection mix was vortexed briefly and incubated at room temperature for 10 min before adding to the cells. Cell medium was refreshed the next day using 7 mL medium, the conditioned medium was discarded. 2 days after transfection, conditioned medium was harvested and stored at 4 °C and 7 mL fresh medium was applied. 3 days after transfection medium was harvested and combined with medium harvested the previous day, spun for 5 min at 3000 rpm to pellet any floating cells, then supernatant was filtered through a 0.045 μm filter to remove debris. Filtrate was applied to an Amicon Ultra-15 filter column (Merck, UFC903029) and centrifuged for 30 min at 3000 rpm to purify virus. Remaining supernatant with virus was aliquoted and stored at -80 °C until used for cell transduction.

HT1080 WT or GEF-H1 KO were plated in a 12 well plate such that they would be 40-50% confluent on the day of transduction. Medium was refreshed with medium containing 6 μg/mL polybrene (Merck, TRI-1003-G) prior to applying virus. Lentivirus was thawed on ice and 5 μL applied per well. 24 hours after transduction medium was refreshed with medium containing 3 μg/mL puromycin (InvivoGen, anti-pr-1) to select for transduced cells. After 24 hours of antibiotic selection cells were returned to normal cell culture medium, expanded and frozen down for future use.

## RhoA2G FRET experiments

RhoA2G HT1080 WT or GEF-H1 KO cells were plated on 25 mm coverslips and serum starved as described above. Just before the start of the experiment, coverslips were mounted inside prewarmed imaging rings (Thermo Scientific, A7816) with 1 mL serum free media. FRET experiments were conducted on a Zeiss LSM880 Airyscan microscope fitted with an S1 (Pecon, Germany) temperature control and CO_2_ system set to 37 °C and 5% CO2, and a 100x/1.46 Alpha Plan-APO oil immersion objective. ZEN black v2.3 software was used to operate the microscope, a 445 nm laser was used to excite the mTFP, the emission for mTFP was collected between 472-500 nm while the emission for mVenus was collected between 565-654 nm; these settings were previously optimized to minimize crosstalk using cells containing only mTFP or mVenus as controls. 1 Frame was acquired every 2 minutes and 5 frames were acquired immediately prior to drug/ dimethyl sulfoxide (DMSO) or serum addition.

## RhoA G-LISA GTPase activation assay

A RhoA G-LISA GTPase activation assay kit purchased from Cytoskeleton (BK121) was used according to manufacturer’s instructions to quantify active RhoA in cells. HT1080 cells were plated at 10% confluency in 10 cm dishes and serum starved as described above. Cells were treated with either DMSO, paclitaxel or docetaxel for the concentrations and times indicated before cells were washed on ice in a refrigerated room (4 °C) with ice cold phosphate-buffered saline (PBS) (taking care to remove all the PBS by aspirating at an angle on ice for 1 minute) and lysed with ice-cold lysis buffer from the kit. Cell lysates were homogenized using a syringe with a 27 gauge needle and pre-cleared by centrifuging in a centrifuge pre-cooled to 4 °C for 2 min at 10,000 rpm. Supernatant was aliquoted on ice into ice cold Eppendorf tubes, snap frozen and stored at – 80 °C until the start of the G-LISA essay. One of the aliquots was used for quantifying the protein concentration using a BCA assay. A final concentration of 250 ng protein / µL was used in the assay after dilution with the binding buffer. ELISA was conducted carefully following manufacturer’s instructions. A FLUOstar OPTIMA plate reader was used to quantify luminescence. Each experiment contained duplicates, the average of the background (lysis buffer only) was subtracted from the average of the sample, and the result was normalized to the mean of the experiment.

## Immunofluorescence cell staining

For staining F-actin, 4% paraformaldehyde (PFA) fixation was used and for staining microtubule binding proteins such as GEF-H1 or EB1 + EB3, methanol fixation was used. For PFA fixation, cells were fixed at room temperature for 15 min using 4% PFA in MRB80 buffer (80 mM K-Pipes, 1 mM EGTA, 4 mM MgCl₂; pH 6.80 with KOH) prewarmed to 37 °C. For methanol fixation, cells were fixed on ice for 15 min using -20 °C methanol. After fixation, cells were washed 3x with PBS. In the case of 4% PFA fixation, cells were permeabilized with 0.2% Triton-X in PBS for 2.5 min. Cells were then blocked in blocking buffer (2% Bovine Serum Albumin in PBS with 0.05% Tween 20) for 1 hour. Primary antibodies were diluted in blocking buffer and incubated on cells for 1 hour, antibodies used were: Rat anti-tyrosinated-α-tubulin (Thermo, MA1-80017; dil 1:200), Mouse anti-α-tubulin (Sigma, T6199; dil 1:200), Mouse anti-acetylated-tubulin (Sigma, T7451; dil 1:200), rabbit anti-de-tyrosinated-tubulin (Abcam, ab48389, dil 1:200), rat EB1 + EB3 (Absea, 15H11; dil 1:10), rabbit anti-GEF-H1 (Abcam, ab155785; dil 1:150), mouse anti GFP (Sigma, 11814460001; dil 1:200). Coverslips were washed by dunking in two different containers of PBS with 0.05% Tween 20, then secondary antibodies or phalloidin diluted in blocking buffer were incubated on cells for 1 hour, secondary antibodies or phalloidin used were: donkey anti-rat-Alexafluor-488 (Thermo, A21208; dil 1:200), goat anti-rat-Alexafluor-594 (Thermo, A11007; dil 1:200), AlexaFluor-594 conjugated phalloidin (Thermo, A12381; dil 1:200), goat anti-mouse-AlexaFluor-488 (Thermo, A11029; dil 1:200) goat anti-mouse-DyLight-405 (Jackson ImmunoResearch, 115-475-166; dil 1:150), goat anti-rabbit-Alexafluor-405-plus (Thermo, A48254; dil 1:200), goat anti-rabbit-AlexaFluor-488-plus (Thermo, A32731; dil 1:200). Samples were mounted with Mowiol mounting medium (10% Mowiol, Sigma, 81381; 30% glycerol, 60% 0.2 M Tris, pH 8.5) with 2.5% DABCO (Sigma, D2522) and allowed to set overnight.

Images of fixed cells for analysis were acquired on a Nikon Exlipse 80i widefield microscope using a Photometrics CoolSNAP MYO CCD camera and a Plan Apo VC 100x 1.40 oil objective. A Chroma ET-DAPI (49000) filter was used to image AlexaFluor-405 and DyLight-405, a Chroma ET-GFP (49002) filter was used to image AlexaFluor-488 or GFP, and a Chroma ET-mCherry (49008) filter was used to image AlexaFluor-594. Nikon NIS Br software was used to operate the microscope.

Images of cells for figure panels in Figure 2 and 3 were acquired on a Leica TCS SP8 confocal microscope using a 100x (oil) HC PL APO 100x/1.40 objective. Microscope is equipped with a 405 nm DMOD Flexible laser and a White Laser (programmable between 470-670 nm), 2 PMT and 3 HyD detectors. LAS X software was used to control the microscope.

## Live cell imaging of GFP-GEF-H1 with drug addition

U2OS seeded on 25 mm glass coverslips were transfected with GFP-GEF-H1 and mCh-α-tubulin as described above. 48 hours after transfection coverslips were mounted in prewarmed imaging rings (Thermo, A7816) with 1 mL medium. Cells were imaged on a custom spinning disc microscope with an Eclipse Ti-E body (Nikon) with perfect focus, a MS-2000-XYZ stage with Piexo Top Plate (ASI), a CSU-X1-A1 spinning disc unit (Yokogawa) and STXG-PLAMX-SETZ21L temperature control and CO_2_ (TokaiHit). S Fluor 100x oil objective (Nikon) and stage were preheated to 37 °C with 5% CO_2_. Voltran Stradus 488 nm (Vortran) and Coherent OBIS 561 nm (Coherent) lasers were used to excite the sample and ET525/50m (GFP) and ET630/75m (mCherry) filters were used to filter signal. A prime BSI sCMOS camera (Teledyne Photometrics) was used to take images, and MetaMorph 7.10 (Molecular Devices) software was used to control the microscope. One image was taken every 30 seconds, the first 10 minutes acquired served as a baseline, after which either 10 µM paclitaxel (PTX) or discodermolide (DDM) was applied to cells and cells were imaged for another 30 minutes.

## Analysis of cell images

### FRET analysis

Fluorescence intensities of mTFP/donor and mVenus/acceptor channels for FRET experiments were quantified in Image J. Cell outlines were made, and the mean pixel intensity (mean grey value) quantified for both channels at all time points, alongside their respective background measurements, taken at an area without cells. FRET ratio 𝑟_𝐹𝑅𝐸𝑇_ was subsequently calculated using the equation below, where 𝐼_𝑑𝑜𝑛𝑜𝑟_ and 𝐼_𝑎𝑐𝑐𝑒𝑝𝑡𝑜𝑟_ are the intensities of the donor and acceptor channels respectively.

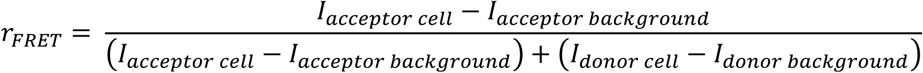

### Quantification of fluorescence intensity

GFP-GEF-H1 fluorescence intensity or Phalloidin staining intensity was measured in Image J by outlining the cell and quantifying the mean pixel intensity (mean grey value) and subtracting the mean pixel intensity of an area without cells (background). Final values were normalized to the experiment’s mean.

### Quantification of GFP-GEF-H1 binding to microtubules

The GFP-GEF-H1 microtubule to cytoplasmic ratio 𝑟_𝑀𝑇/𝑐𝑦𝑡𝑜𝑝𝑙𝑎𝑠𝑚_ was quantified in Image J. First, channels were aligned using the template matching plugin. Next, mean GFP-GEF-H1 fluorescence was determined for all cells as described above and only cells with comparable expression levels across different treatment groups were used for subsequent analysis. For each cell, 10 line scans, 3 pixels wide, were drawn perpendicular to the microtubule in regions where the cytoplasm was devoid of microtubules on either side of the microtubule. The GFP-GEF-H1 intensity on the microtubule 𝐼_𝑀𝑇_ was calculated by taking the mean pixel intensity at the location with the maximum value of the tubulin staining plus the value directly on either side of that maximum; the GFP-GEF-H1 intensity in the cytoplasm 𝐼_𝑐𝑦𝑡𝑜𝑝𝑙𝑎𝑠𝑚_ was defined as the values 4-6 pixels before and after the maximum of the tubulin staining. These distances were determined by looking at line scans across microtubules as shown in Fig. 3B. The mean background pixel intensity 𝐼_𝑏𝑎𝑐𝑘𝑔𝑟𝑜𝑢𝑛𝑑_ was measured at a region devoid of cells. The background was subtracted from the GFP-GEF-H1 intensity on the microtubule and in the cytoplasm, and then the ratio was taken of GFP-GEF-H1 on the microtubule and GFP-GEF-H1 in the cytoplasm as follows:

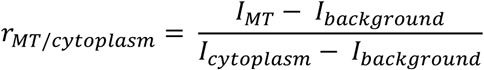

The mean cell microtubule to cytoplasm ratio over the 10 line scans was plotted. GEF-H1 microtubule to cytoplasm ratio was analyzed for 11-12 cells per treatment group per experiment.

## Protein Purification

The proteins were purified from HEK293T cells using the Strep(II)-streptactin affinity purification as described previously (Saunders et al., 2025). Cells were transfected with a 1:3 w/w mix of DNA construct and PEI (Polysciences) and harvested 48h after transfection in lysis buffer (50 mM HEPES, 1 mM MgCl_2_, 1 mM DTT, 300 mM NaCl, and 0.5% Triton X-100; pH 7.4) containing protease inhibitors (Roche). They were kept on ice for 15min and centrifuged to clear the debris. The lysate was incubated with StrepTactin beads (IBA lifesciences) for 45mins. The beads were washed five times with lysis buffer (high salt washes) and three times with lysis buffer but with 150mM NaCl instead of 300mM and (low salt washes). The protein was then eluted in elution buffer (50 mM HEPES, 150 mM NaCl, 1 mM MgCl_2_, 1 mM EGTA, 1 mM DTT, 0.05% Triton X-100 and 2.5 mM d-desthiobiotin (Sigma-Aldrich), pH 7.4). All purified proteins were snap-frozen and stored at -80°C.

## Mass Spectrometry

### Protein digestion

For liquid chromatography-tandem mass spectrometry (LC-MS/MS) analysis, 2 mg GFP-GEF-H1 purified protein was used. The volume was adjusted to 20 µL using elution buffer [50 mM HEPES pH 7.4, 150 mM NaCl, 1 mM MgCl_2_, 1 mM EGTA, 1 mM DTT, 0.05% Triton X-100, 2.5 mM d-Desthiobiotin]. The sample was first reduced and alkylated by adding Tris(2-carboxyethyl)phosphine (TCEP) and chloroacetamide (CAA) to a final concentration of about 10 mM and 40 mM, respectively. Incubation first took place at 99 °C for 10 minutes, then in the dark for 20 minutes at room temperature (RT).

The sample was digested using the single-pot, solid-phase-enhance sample preparation method (SP3) (Hughes et al., 2019). For this, the Cytiva Sera-Mag™ Carboxylate-Modified Magnetic Beads, both hydrophilic and hydrophobic beads were used in a 1:1 ratio. The required amount of beads was first washed with ultrapure water (MQ), which was repeated twice. To make sure the bead concentration remained above 0.5 µg/uL, a total amount of 75 µg beads were added to the sample. 100% ethanol (EtOH) was added to the sample to a final concentration of 75% EtOH. The sample was incubated in a Thermomixer at 1 000 rpm for 20 minutes at RT. The tube was placed in a magnetic rack, and the supernatant was removed. The beads were washed two times with 80% EtOH, followed by a washing step with 100% acetonitrile (ACN). For digestion, the beads were resuspended in 200 µL 100 mM ammonium bicarbonate (AMBIC) by pushing the particles into solution; the digestion-bead solution was not pipetted or homogenized. Then, the sample was sonicated for two minutes in a water bath and digested by using the proteases trypsin and lys-C, in a 1:25 and 1:75 protease to protein ratio, respectively. Digestion was performed overnight in a Thermomixer at 1 000 rpm, at 37°C. The sample was spun down and acidified by adding 10% trifluoracetic acid (TFA) to a final concentration of 5% TFA. The beads were immobilized on the magnetic rack, and the acidified peptide solution was transferred to a new Eppendorf tube. The peptide sample was dried in a vacuum centrifuge.

## Liquid Chromatography with tandem mass spectrometry (LC-MS/MS)

The sample was resuspended in 50 µL 2% formic acid (FA), and 5 µL was used to analyze with LC-MS/MS. Peptides were separated on the nanospray UHPLC system Ultimate 3000 (Thermo), using a 50-cm reversed-phase analytical column, with an integrated emitter, that has a 75 µm inner diameter, and was packed in-house with ReproSilPur C18- AQ 1.9 µm resin (Dr. Maisch GmbH). Mobile Phase Buffer A was composed of 0.1 % FA (v/v) and Mobile Phase Buffer B consisted of 80% ACN (v/v) and 0.1% FA (v/v). The Acclaim Pepmap 100 C18 (5 mm × 0.3 mm, 5 μm, Thermo) trap column was operated at a temperature of 32°C, and the analytical column at a temperature of 50°C. A flow rate of 0.3 μL/min was used. Starting at 4.0%, Buffer B was increased over a total gradient-time of 45 minutes using the following stepwise increases: 11% Buffer B (3-35 min), 30% Buffer B (35-40 min), and 44% Buffer B (40-44 min), and 55% Buffer B (44-45 min). The column was then washed out with 99% Buffer B for 5 minutes, followed by a 10-minute re-equilibration step with 4.0% Buffer B.

The Ultimate 3000 was coupled to a Thermo Scientific™ Orbitrap Exploris 480. Data was acquired in a data-dependent mode (DDA), and MS1 scans were acquired at a 60 000 resolution using a scan range of 375-1600 m/z. With an isolation window of 1.4 m/z, precursor ions with an intensity threshold higher than 200 000, and with charge states between 2+ and 6+ were selected for fragmentation. Precursors were fragmented by stepped higher-energy collisional dissociation (HCD) with a Normalized Collision Energy (NCE) of 28%. The MS2 scans were acquired at a 15 000 resolution, with maximum injection time set to auto, and a scan range set to a first mass of 120 m/z. Dynamic exclusion time was set to custom, with a 10-p.p.m. upper- and lower tolerance and excluding isotypes.

## MaxQuant Analysis

The raw data was analyzed using the MaxQuant-Andromeda software (v.2.4.7.0) for protein identification, using the iBaq values for quantification. The MS/MS spectra were searched against the UniProtKB Human Proteome (organism_id: 9606, reviewed, canonical & isoform FASTA, downloaded on: 12th of March, 2025), and a separate FASTA file containing the recombinant protein, StrepII-GFP-GEF-H1. Default parameters were used for the precursor mass tolerance (20 p.p.m first search, 4.5 p.p.m. main search), and the False-Discovery Rate (FDR) was set to 1%. Up to three missed cleavages for both proteases were allowed. Carbamidomethylation of cysteine residues was set as a fixed modification, and oxidation of methionine and acetylation of the protein N-terminus were set as variable modifications, with a maximum of five modifications per peptide.

The iBaq values from the MaxQuant search were imported into the Perseus software (v.2.0.11.0), where the protein groups were filtered out for ‘Reverse’ (false positives), ‘Contaminants’, and ‘Only identified by site’, as well as having only one peptide hit, and transferred to excel files for further analysis. The iBaq values were normalized to percentages, were the highest iBaq value was set to 100%.

## *In vitro* reconstitution assays Sample preparation

Flow chambers were prepared by plasma cleaning coverslips and attaching them to glass slides using strips of double-sided tape to create ∼10 µL volume chambers. The surface was functionalized by 5-minute incubations with 0.2 mg/mL PLL-PEG-Biotin (Susos AG) and then 0.83 mg/mL neutravidin (Invitrogen), both dissolved in MRB80 buffer (80 mM K-Pipes, 1 mM EGTA, 4 mM MgCl₂; pH 6.80 with KOH). Double-cycled biotinylated GMPCPP seeds prepared as described previously (van den Berg et al., 2023) were then flowed in and incubated for 2 minutes, followed by a >3 minute incubation with κ-casein.

All tubulin reagents used were from Cytoskeleton Inc. All mixes contained a freshly prepared master mix composed of 0.1% (w/v) methylcellulose, 0.5 mg/ml κ-casein, 50 mM KCL, 50 mM glucose, 0.2 mg/ml catalase, 0.5 mg/mL glucose oxidase, and 10 mM DTT, and were mixed in the MRB80 buffer. For the GFP-GEF-H1 binding assays, the master mix was supplemented with 17 µM porcine brain tubulin, 0.6 µM TRITC rhodamine labelled tubulin, 1 mM GTP/GTPyS and the required concentration of GFP-GEF-H1 proteins. For the paclitaxel and discodermolide expansion assays, the microtubule mix had master mix supplemented with 17 µM porcine brain tubulin, 0.6 µM TRITC rhodamine labelled tubulin and 1 mM GTP; and the wash-in mix had master mix with 20 µM paclitaxel or 200 nM discodermolide. For the experiments with GEF-H1 and paclitaxel or discodermolide, the microtubule mix was the same as the binding assays and the wash-in mix had 4.85 µM porcine brain tubulin, 0.15 µM TRITC rhodamine labelled tubulin, 20 µM paclitaxel or 200 nM discodermolide and the same concentration of GEF-H1 proteins. All mixes were spun in an Airfuge for 5 min at 119,000 x g. The microtubule mixes were flowed into the chambers, and the wash-in mixes were kept on ice before manually washing-in during image acquisition.

### TIRF microscopy

Total internal reflection fluorescence (TIRF) microscopy was performed on an inverted Nikon Eclipse Ti-E research microscope (Nikon) equipped with azimuthal spinning TIRF illumination, a perfect focus system, and an ASI motorized MS-2000-XY stage. Imaging was carried out using a Nikon APO TIRF 100× 1.49 NA oil immersion objective in combination with an iLas2 illumination system (Roper Scientific, now Gataca Systems). Images were acquired with a CoolSNAP MYO CCD camera (Teledyne Photometrics) providing a pixel size of 0.045 μm/pixel. Samples were maintained at 30 °C for all experiments using a Tokai Hit STXG-PLAMX-SETZ21L Stage Top Incubator. Excitation was achieved using the following lasers: Coherent OBIS 561 nm (100 mW), Vortran Stradus 642 nm (110 mW), and Vortran Stradus 488 nm (150 mW). Emission was collected through the corresponding Chroma filters ET-GFP (49002), ET-mCherry (49008), and ET-647. Image acquisition was controlled using Metamorph software (version 7.10.2.240, Molecular Devices).

### Image processing

To calculate the intensity, the movies were averaged over frames, and the mean intensity was calculated at the required region of interest. The ImageJ plugin KymoResliceWide v.04 was used to generate Kymographs (https://github.com/ekatrukha/KymoResliceWide). To calculate lattice expansion, a straight line was drawn from the end of the bright seed to a speckle toward the plus end of the kymograph before the wash-in. After the flow-in, a second line was drawn between the same two points. The difference in lengths, reflecting the speckle shift, was used to calculate the percentage of expansion.

## Interference Reflection Microscopy (IRM) to determine microtubule protofilament number

Porcine brain tubulin was isolated using the high-molarity PIPES method (Castoldi and Popov, 2003). Biotin-labeled tubulin (Cytoskeleton Inc, T333P) was diluted 1:50 with unlabeled porcine brain tubulin to obtain biotin-labeled tubulin mix. 12 protofilament microtubules were polymerized from 4 mg/ml biotin-labeled tubulin for 1 hour at 37°C in 10 mM phosphate buffer supplemented with 6 mM MgCl_2_, 2 μM paclitaxel, and 1mM GTP (NU-1012, Jena Bioscience) according to (Ray et al., 1993). Polymerized microtubules were centrifuged for 30 min at 18000 x g in a Microfuge 18 Centrifuge (Beckman Coulter). After centrifugation the pellet was resuspended and kept in BRB80 supplemented with 10 μM paclitaxel at room temperature. GMPCPP-stabilized 14 protofilament microtubules (and seeds for dynamic microtubule assay) were polymerized from 4 mg/ml biotin-labeled tubulin for at least 3 hours at 37 °C in BRB80 supplemented with 1 mM MgCl_2_ and 1 mM GMPCPP (Jena Bioscience, NU-405). Polymerized microtubules were centrifuged, and the pellet was resuspended as above.

For imaging experiments, chambers were assembled by melting thin strips of parafilm in between two glass coverslips silanized with Hexamethyldisilazane (HMDS, 379212) and functionalized with 20 µg/mL anti-biotin antibodies (in BRB80; Sigma, B3640) incubated for 5 min and 1% Pluronic (F127 in BRB80; Sigma, P2443) incubated for at least 30 min.

To measure the IRM signal of microtubules, first the background signal was measured, then the 12 protofilament microtubules were immobilized in the chamber and their position was imaged. Then the 14 protofilament GMPCPP stabilized microtubules were added to the chamber and microtubules were imaged. Dynamic microtubule assay was performed by immobilizing GMPCPP seeds on the coverslip surface. Then 8 mg/ml of free fluorescent tubulin HiLyte488 (Cytoskeleton Inc, TL488M) was added in the BRB80 based assay buffer (10 mM DTT, 20 mM D-Glucose, 0.1% Tween 20, 0.05 mg/ml Casein, 1 mM ATP, 1 mM GTP, Glucose oxidase and Catalase). Microtubule dynamics was measured for 5 min using both IRM and TIRF microcopy. IRM signal of the microtubules was measured from the last frame.

Background-subtracted IRM signal, proportional to local protein density, was used to analyze microtubule thickness (protofilament number (Mahamdeh and Howard, 2019). Then the mean IRM signal was measured by drawing a thick line along the microtubule. The resulting number was normalized to the signal of 14 protofilament microtubules for both 12 vs 14 protofilament experiment as well as for the dynamic assay experiment.

## Cryo-Electron microscopy (Cryo-EM) Sample preparation

To ensure tubulin was fully depolymerized, after thawing 50 µM porcine tubulin (Cytoskeleton) was incubated with 2 mM GTP in MRB80 on ice for 5 mins. To reduce the formation of sheet-like structures, this mix was first polymerized for 10 mins at 37 °C in the absence of discodermolide, followed by further polymerization for 20 mins at 37 °C in the presence of 100 µM discodermolide. The polymerized microtubules were then pelleted with the Airfuge® Air-Driven ultracentrifuge (5 min, 119,000 x g). The pellet was resuspended in MRB80 supplemented with 100 µM discodermolide to an approximate concentration of 8 µM tubulin (assuming 80% loss). The concentration of polymerised tubulin was determined by depolymerisation of a sample in MRB80 with 50 mM CaCl_2_ and 1 mM DTT, and measurement at A280 (extinction coefficient: 115000 M^-1^cm^-1^) on a NanoDrop. Finally, discodermolide-bound microtubules were diluted to a concentration of 2 µM tubulin in MRB80 + 100 µM discodermolide.

## Cryo-EM grid preparation and data acquisition

QUANTIFOIL® Holey Carbon R3.5/1 Cu 200 mesh grids (Quantifoil Micro Tools GmbH) were glow discharged at 25 mA for 45 s at 0.39 mbar (PELCO easiGlow™). 4 µL of discodermolide stabilised microtubules were applied to the grid in the EM GP2 Automatic Plunge Freezer (Leica) chamber. Grids were incubated at 30 °C and 95 % humidity for 30 s, back-side blotted for 1.5 s and then plunged into liquid ethane. Data was recorded on a Titan Krios I (The Netherlands Centre for Electron Nanoscopy (NeCEN), Leiden University, The Netherlands) equipped with a K3 summit direct electron detector and operated at 300 keV using EPU. A magnification of 105000x corresponding to a pixel size 0.836 Å at the sample was used, with a total dose of 50 e-/Å^2^ per exposure collected over 50 fractions. A defocus range of -0.6 to -2.0 µm was used.

## Cryo-EM image processing

Movies were motion corrected and CTF estimation was performed using cryoSPARC (Punjani et al., 2017; Rubinstein and Brubaker, 2015). Approximately 500 particles from movies with varied defocus values and CTF estimations were manually picked, 2D classified and used as input for automatic filament tracing in cryoSPARC (Punjani et al., 2017; Rubinstein and Brubaker, 2015). Filament tracing was performed with the separation distance set to the length of a single dimer 82 Å. Particles were extracted in a box size twice as big as microtubule filament diameter (∼500 Å, 576 pixels) and binned by 2 (288 pixels). The particles were sorted into 11- to 16-protofilament classes by heterogeneous refinement. Particles belonging to 13 PF microtubules and 14 PF microtubules were selected for further separate processing. First, helical refinement was performed (twist = 0 °, rise = 82.5 Å), followed by local refinement. Unbinned particles were then re-extracted in cryoSPARC, (Punjani et al., 2017; Rubinstein and Brubaker, 2015) and refined without any imposed symmetry (C1). Lattice parameters were calculated from a helical search of the C1 map using relion helix toolbox in Relion 5.0 (He and Scheres, 2017).

## Statistics

The Kolmogorov-Smirnov test was performed to determine whether the data fit a normal distribution for >4 data points and the Shapiro Wilk test was used for 3-4 data points. The F-actin intensity data, cell circularity data, IRM data and all the *in vitro* data was determined not to fit a normal distribution and therefore a Mann-Whitney U test was used to test for significance between treatment and control groups. Data on acetylated and non-acetylated GEF-H1 microtubule to cytoplasmic ratio was also determined not to follow a normal distribution. In this case, a Wilcoxon matched-pairs signed rank test was used, since both GEF-H1 binding on acetylated and un-acetylated microtubules was quantified in each cell. The GTP-RhoA ELISA data, p-MLC blot quantifications, GEF-H1 microtubule to cytoplasm ratio data and GFP-GEF-H1 intensity data were determined to follow a normal distribution, and therefore a Welch’s t-test that does not assume equal variance was employed to test for significance between treatment and control groups. A P value below 0.05 was determined to be significant for all of the above-mentioned tests.

## Data Availability

All other data that support conclusions of this paper are either available in the manuscript itself or available from the authors on request. All unique and stable reagents generated in this study are available from the lead contact without restriction.

---

**Figure S1.**
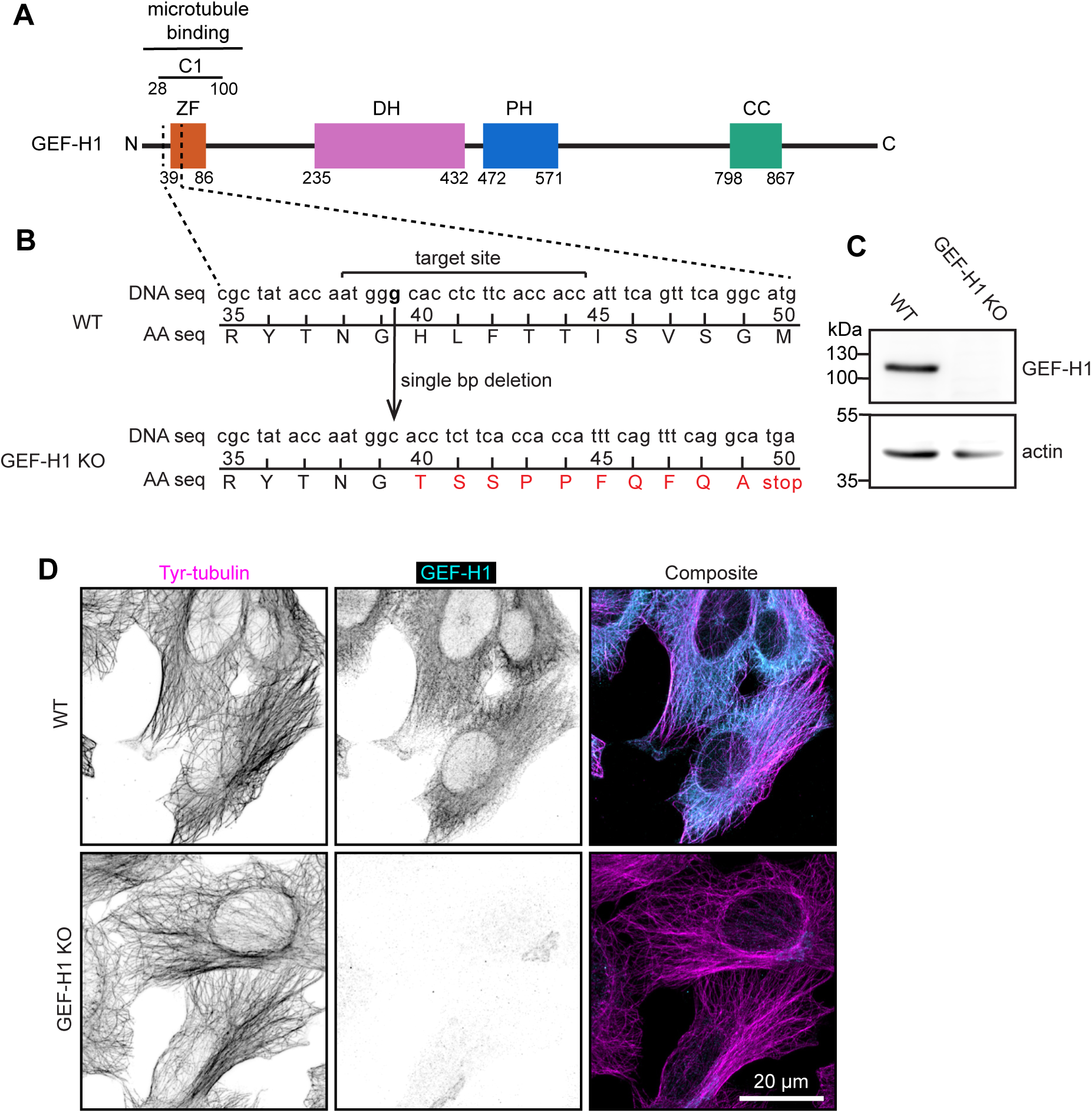
**Characterization of HT1080 GEF-H1 knockout (KO) cell line.** (A) Map of GEF-H1 protein domains, showing Zinc Finger (ZF), Dbl homology domain (DH), Pleckstrin homology domain (PH), Coiled Coil (CC) adapted from the Uniprot database, and the C1 microtubule binding domain described in (Choi et al., 2025). (B) Scheme showing the location of the single DNA base pair (bp) deletion in the GEF-H1 gene of GEF-H1 KO cells, verified by sequencing. (C) Western blot of WT and GEF-H1 KO HT1080 lysates blotted for GEF-H1 and actin as a loading control. (D) WT and GEF-H1 KO HT1080 cells fixed and stained for GEF-H1 and tyrosinated α-tubulin.

**Figure S2.**
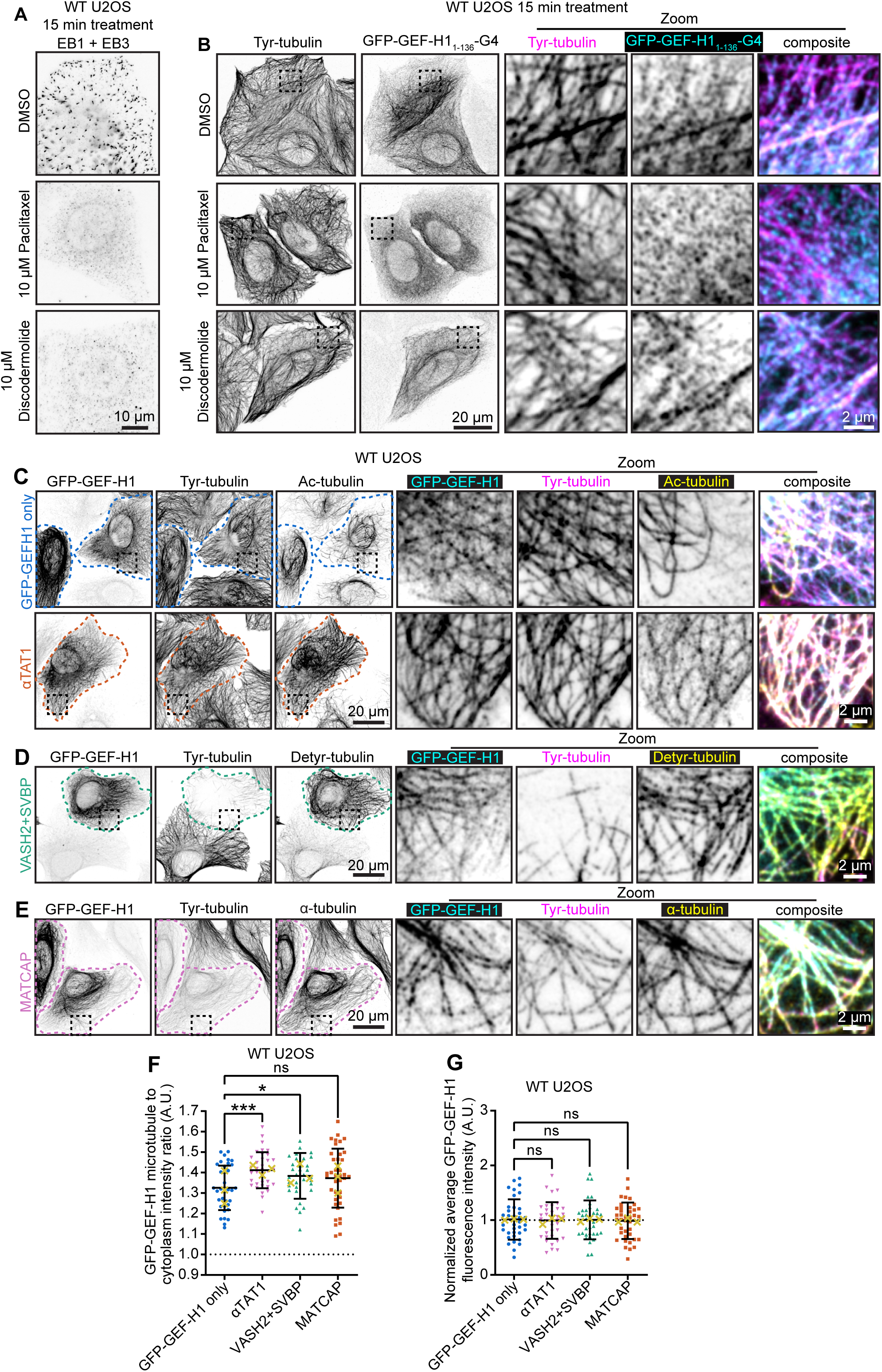
Characterization of cellular effects of MTAs and treatments affecting post-translational microtubule modifications. (A) U2OS cells were treated for 15 minutes with either DMSO, paclitaxel or discodermolide, fixed and stained with antibodies recognizing End Binding proteins EB1 and EB3, which visualize growing microtubule ends. (B) U2OS cells transiently transfected with GFP-GEF-H1_1-136_-G4 were treated for 15 minutes with either DMSO, paclitaxel or discodermolide, fixed and stained for tyrosinated tubulin. (C-G) U2OS stably overexpressing GFP-GEF-H1 at low levels were transiently transfected with αTAT1 (C) to induce acetylation, VASH2 + SVBP (D) to induce detyrosination or MATCAP (E) to induce removal of two C-terminal amino acids of α-tubulin (Δ2). Cells were fixed and stained for tyrosinated tubulin (tyr-tubulin) and either acetylated tubulin (C, ac-tubulin), detyrosinated tubulin (D, detyr-tubulin) or Δ2-tubulin (E). (C-E) Representative confocal images are shown, where colored stippled lines mark transfected cells and dashed boxes indicate which regions are enlarged in the zoom panels. (F) Plot shows GFP-GEF-H1 microtubule to cytoplasm intensity ratio, analyzed in untransfected or αTAT1 / VASH2+SVBP / MATCAP transfected cells as shown in Fig. 3B. (G) Plot shows GFP-GEF-H1 intensity normalized to experimental mean of cells for which microtubule binding was analyzed. (F, G) Individual cell means are shown as colored symbols, yellow crosses show experimental means and brackets show standard deviation based on individual cell values (GFP-GEF-H1 only n = 36, αTAT1 n = 36, VASH2+SVBP n = 35, MATCAP n = 39; from 3 experiments) A Welch’s t-test was used to test for statistical significance, ns = not significant, * = p<0.05, *** = p<0.001.

**Figure S3.**
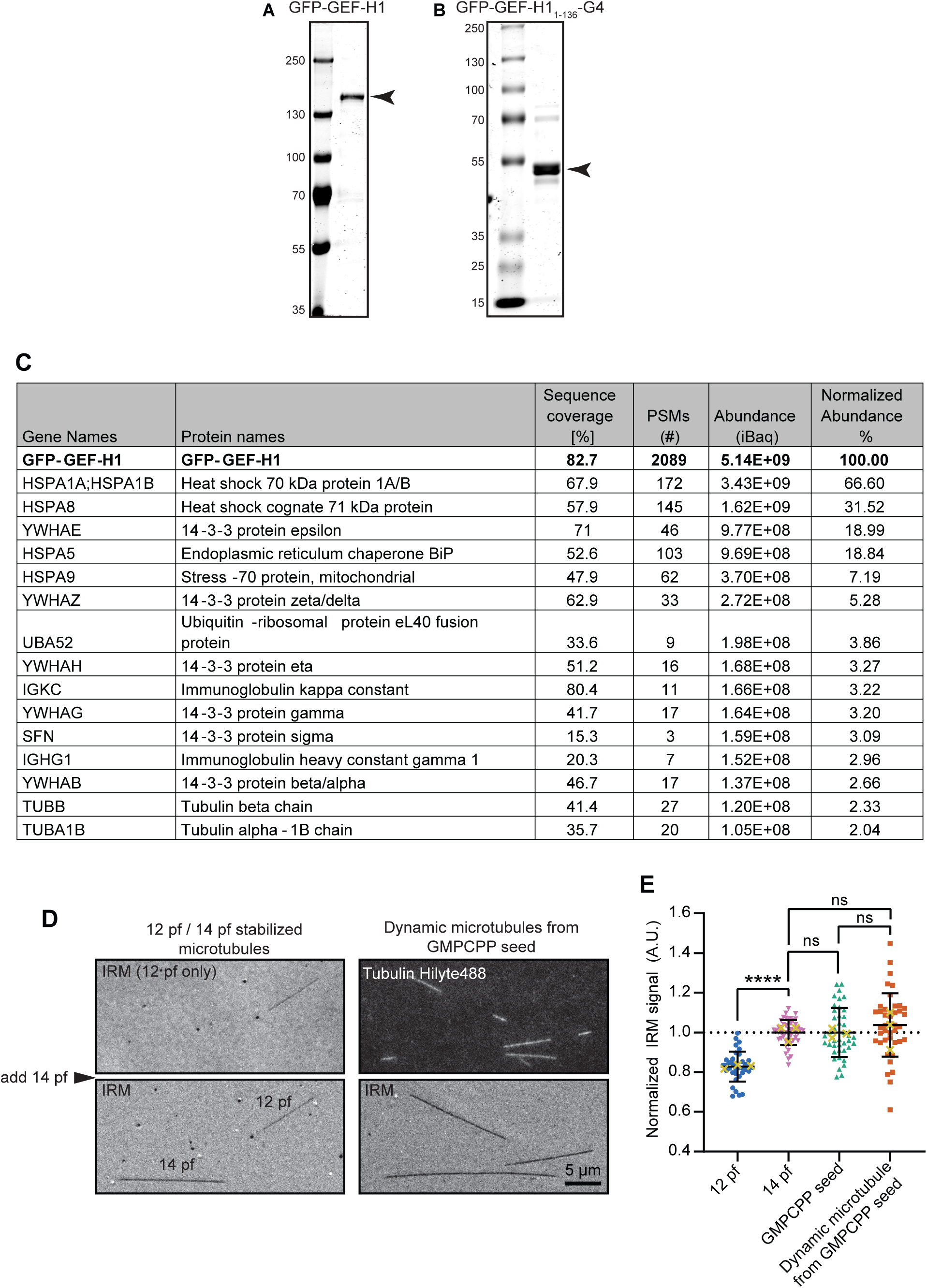
Purified GEF-H1 protein controls and IRM analysis to determine protofilament number. (A, B) Coomassie gels of purified (A) full length GFP-GEF-H1 protein and (B) GFP-GEF-H1_1-136_-G4 protein. (C) Mass spectrometry analysis of purified GFP-GEF-H1 protein. (D, E) Interference Reflection Microscopy (IRM) was used to compare microtubules of known pf number and to compare dynamic GDP-microtubules polymerized from 14 pf GMPCPP microtubule seeds. (E) IRM signal was normalized to the average signal for GMPCPP 14 pf microtubules; plot shows mean ± standard deviation, with individual microtubules analyzed as colored points and experimental averages as yellow crosses (12 pf n = 36, 14 pf n = 47, GMPCPP n = 41, dynamic microtubule n = 41; from 3 experiments). A Welch’s t-test was used to test for statistical significance, ns = not significant, **** = p<0.0001.

**Figure S4.**
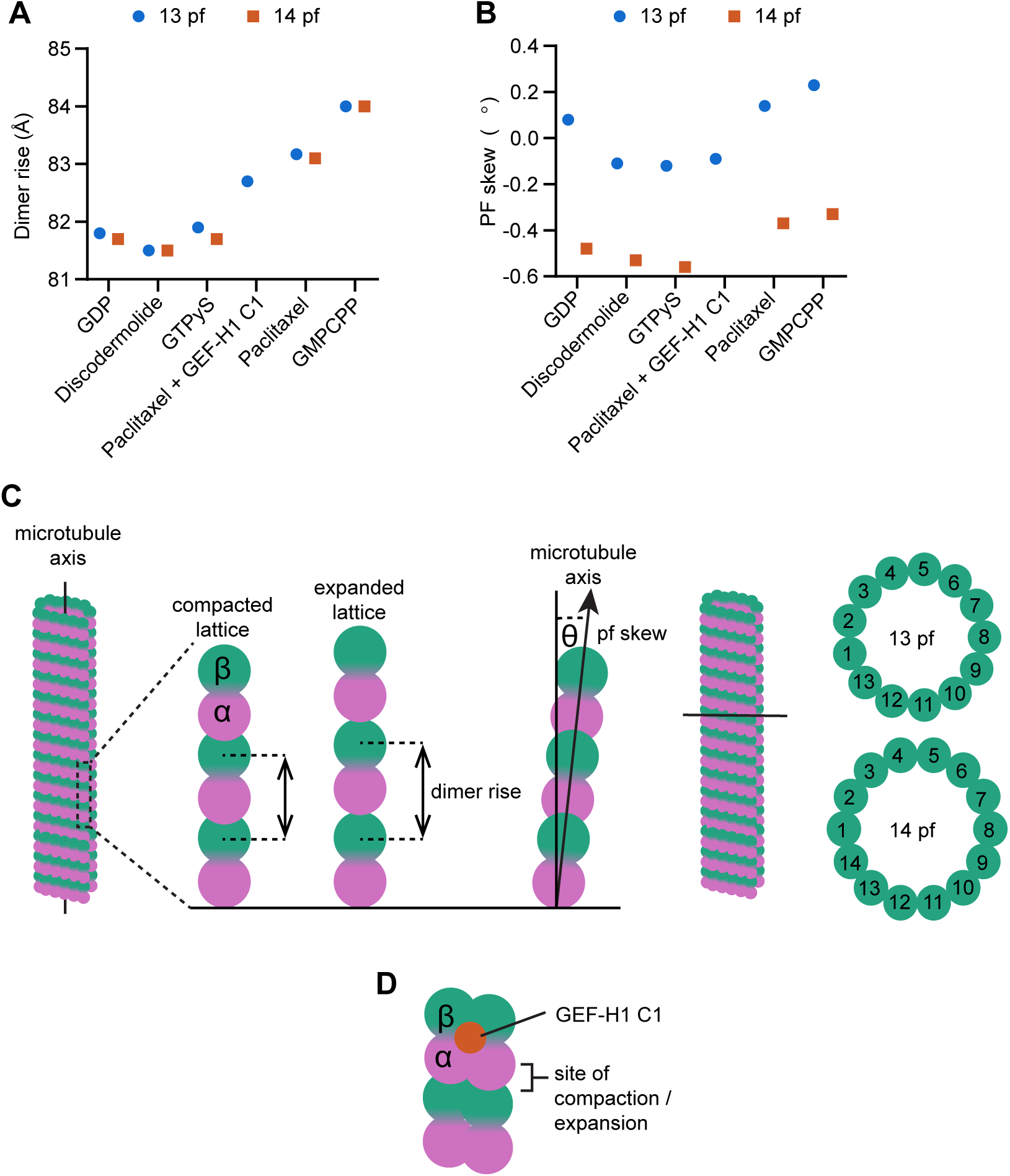
Comparison of microtubule lattice parameters (A, B) Plot of dimer rise (A) and protofilament (pf) skew (B) for 13 pf and 14 pf microtubules with either dynamic GDP lattice or in the presence of discodermolide, GTPyS, paclitaxel + GEFH1-C1, paclitaxel alone or GMPCPP. Values for GMPCPP, GTPyS and GDP lattices are adopted from (Zhang et al., 2018), paclitaxel adopted from (Debs et al., 2020), and paclitaxel + GEF-H1-C1 adopted from (Choi et al., 2025). (C) Illustration of microtubule lattice parameters: dimer rise, which reflects lattice expansion and compaction, pf number and pf skew. (D) Schematic representation of GEF-H1 C1 binding to the microtubule lattice, based on (Choi et al., 2025).

**Figure S5.**
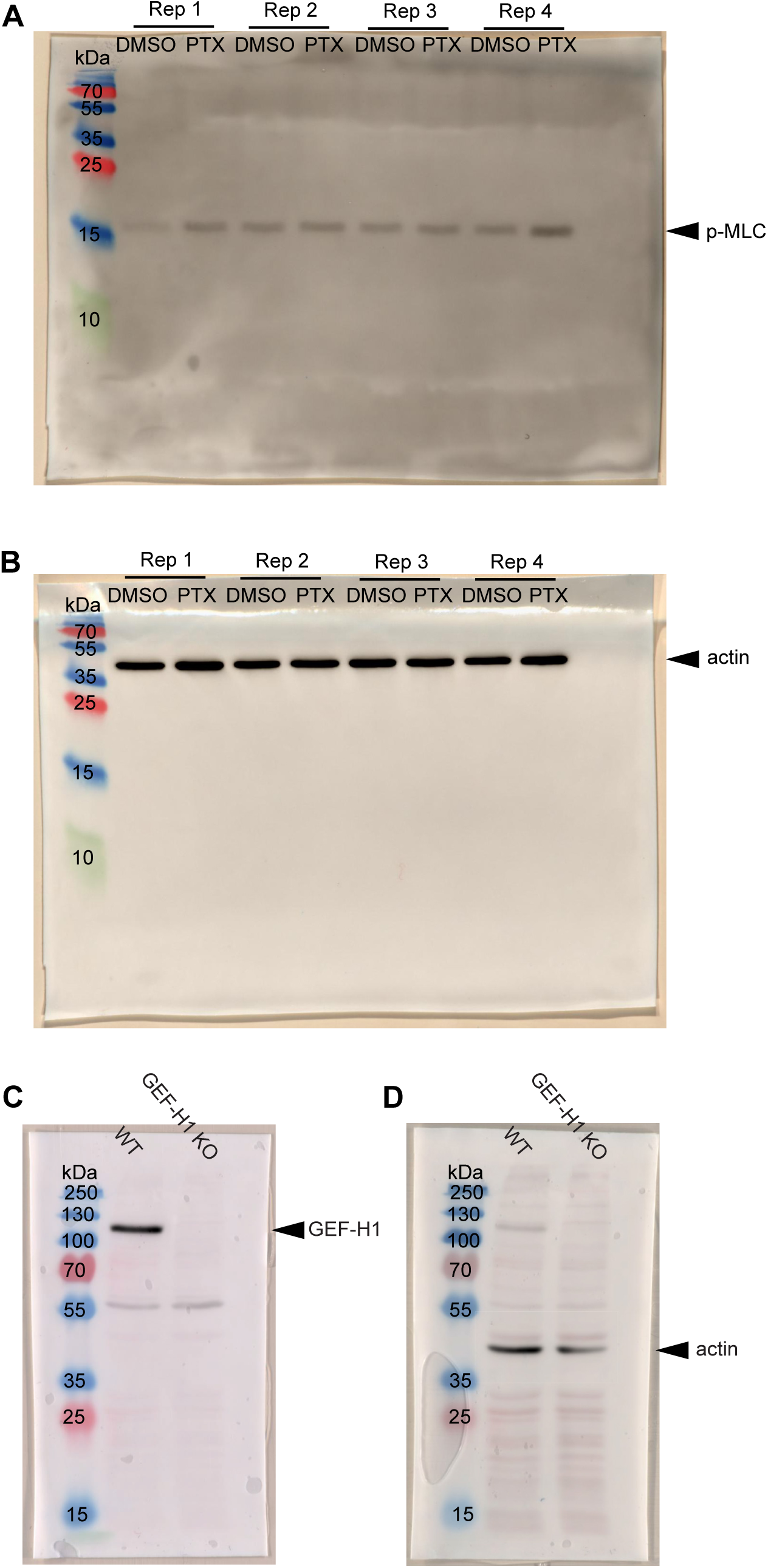
Full Western blots for p-MLC and GEF-H1. (A, B) Serum starved HT1080 WT cells treated with paclitaxel (PTX) or DMSO as a solvent control for 30 minutes before harvesting cell lysates and blotting for (A) p-MLC and (B) actin as a loading control. (C, D) HT1080 WT and HT1080 GEF-H1 KO cells were blotted for (C) GEF-H1 and (D) actin as a loading control.

## Notes

### Competing Interest Statement

The authors have declared no competing interest.

